# The ER Thioredoxin-Related Transmembrane Protein TMX2 Controls Redox-Mediated Tethering of ER-Mitochondria Contacts (ERMCS)

**DOI:** 10.1101/2024.04.12.589228

**Authors:** Junsheng Chen, Megan C. Yap, Arthur Bassot, Danielle M. Pascual, Tadashi Makio, Jannik Zimmermann, Heather Mast, Rakesh Bhat, Samuel G. Fleury, Yuxiang Fan, Adriana Zardini Buzatto, Jack Moore, Klaus Ballanyi, Liang Li, Michael Overduin, M. Joanne Lemieux, Hélène Lemieux, Wen-Hann Tan, Grazia M.S. Mancini, Bruce Morgan, Paul C. Marcogliese, Thomas Simmen

## Abstract

Thioredoxin-related transmembrane proteins (TMX) of the endoplasmic reticulum (ER) have emerged as key regulators of ER membrane properties. Within the ER lumen, TMX proteins and other ER redox enzymes determine oxidative conditions, which control the formation of ER-mitochondria membrane contacts (ERMCS) and determine their function. ERMCS exhibit cytoplasmic redox nanodomains, derived from ER and mitochondrial reactive oxygen species (ROS), whose mechanistic regulation is uncharacterized. Our research has identified the ER protein TMX2, which uses its unique cytosolic thioredoxin domain to prevent cytosolic sulfenylation of mitochondrial outer membrane proteins such as TOM70 through a functional interaction with peroxiredoxin-1 (PRDX1). By doing so, TMX2 interferes with the TOM70 ERMCS tethering function and reduces mitochondrial Ca^2+^ flux and metabolism. Recently, TMX2 mutations have been identified to cause a neurodevelopmental disorder with microcephaly, cortical malformations, and spasticity (NEDMCMS). Using TMX2-mutated NEDMCMS patient cells, we demonstrate that compromising TMX2 through mutation reproduces mitochondrial defects. In a fly *in vivo* model, TMX2 knockdown manifests predominantly in glial cells. Our results therefore provide important mechanistic insight into NEDMCMS and mechanistically link TMX2-mediated control of ERMCS to brain development and function.

**Graphical Abstract:** 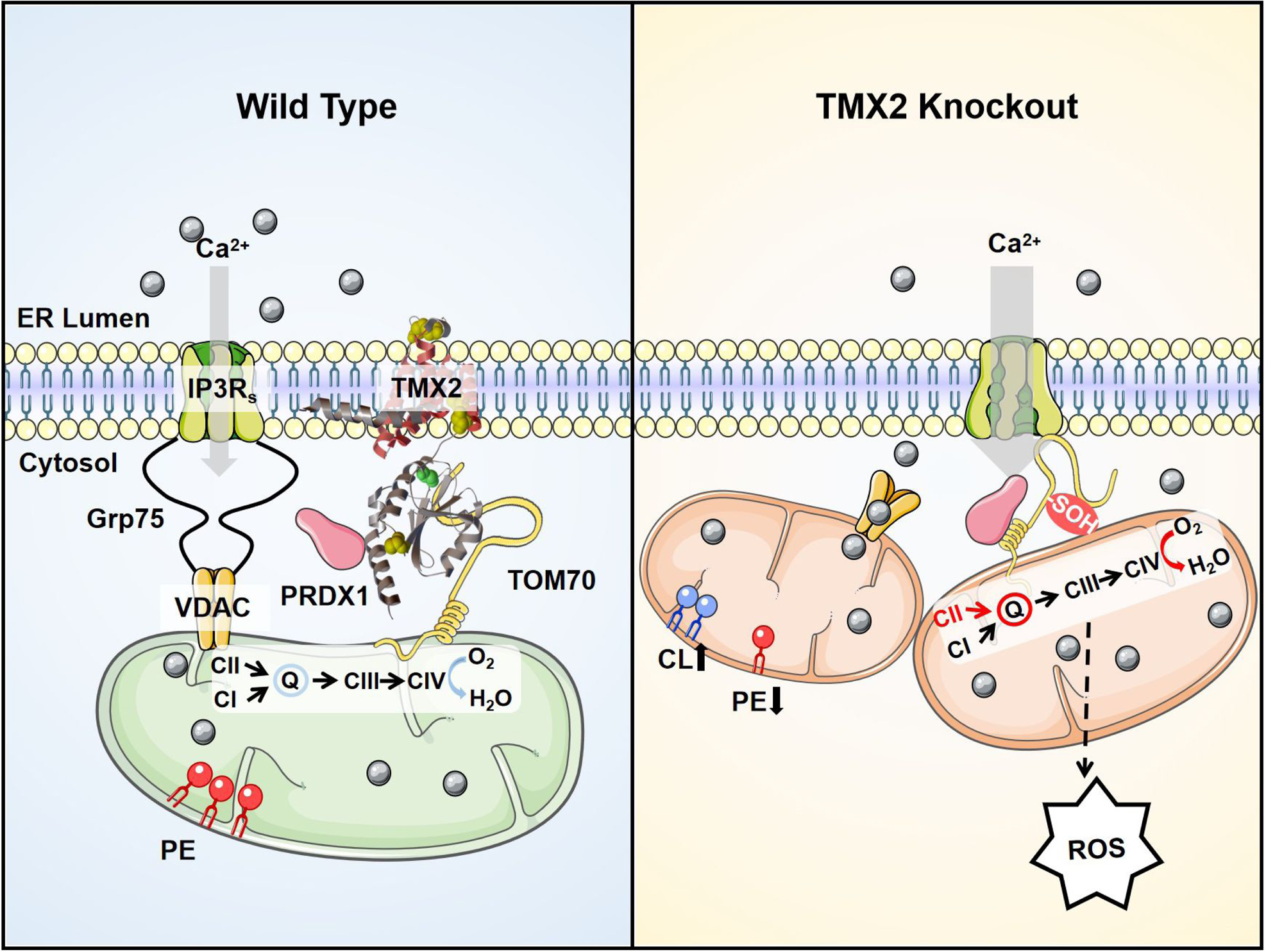

The transmembrane thioredoxin-related TMX2 prevents TOM70 sulfenylation at ERMCS, thus maintaining normal mitochondria metabolism in wild-type cells. TMX2 knockout leads to TOM70 sulfenylation and tight ERMCS formation. This then increases ROS production, unbalances mitochondrial lipids, and relatively shifts OXPHOS electron supply to complex II.

## Introduction

The membranes of the endoplasmic reticulum (ER) contain thioredoxin-related transmembrane proteins (TMX) (Guerra & Molinari, 2020), which can communicate the status of the ER to adjacent organelles, including the transmission of ER stress at the nuclear envelope (Kucinska et al., 2023) or at ER-mitochondria contacts (ERMCS) (Raturi et al., 2016). Importantly, reactive oxygen species (ROS) control ERMCS, reflected in their rich content of multiple ROS producers (Yoboue et al., 2018). This accumulation of H_2_O_2_ can originate from within the ER (Gorin et al., 2003) or from mitochondria (Sharma et al., 2011). An important H_2_O_2_-mediated mechanism operating at ERMCS is based on the multipass transmembrane NADPH oxidase 4 (NOX4), which promotes ER membrane protein oxidation (Beretta et al., 2020), and increases ERMCS Ca^2+^ flux to activate mitochondria (Booth et al., 2021). Similarly, ER oxidoreductin 1 (ERO1) localizes to ERMCS (Gilady et al., 2010), where it produces H_2_O_2_ during the early phases of ER stress. This activity leads to the lumenal oxidation and oligomerization of the UPR sensor protein PERK, increases ERMCS formation, and activates mitochondria (Bassot et al., 2023). These findings suggest that the amount of ROS at ERMCS may determine their function but could also dictate the proportion of tight (<10 nm membrane distance) versus wide ERMCS spacing (>30 nm membrane distance). Such a switch of ERMCS spacing could hypothetically favor lipid or Ca^2+^ signaling, respectively (Giacomello & Pellegrini, 2016).

In contrast to ER lumenal redox control of ERMCS, no cytosolic oxidase or reductase activity has been associated with this structure. In principle, enzymes of the peroxiredoxin family could control cytosolic redox at ERMCS, since they are highly efficient at reducing H_2_O_2_ levels through their peroxidatic and resolving cysteines (Rhee, 2016). This activity extends to the periphery of mitochondria, since peroxiredoxin-1 (PRDX1), a member of the 2-cysteine peroxiredoxin family, absorbs mitochondrial H_2_O_2_ output (Hoehne et al., 2022). Once oxidized, cytosolic thioredoxins reduce PRDXs (Pannala & Dash, 2015). Alternatively, 2-cysteine PRDXs can oxidize substrates in a redox relay, such as towards ASK1 in the case of PRDX1 (Jarvis et al., 2012). The synthesis of these redox reactions can lead to sharp H_2_O_2_ gradients near sources of ROS, generating a localized, PRDX-controlled ROS floodgate (Wood et al., 2003). However, not many cytosolic ERMCS oxidation targets are currently known. One candidate is the ERMCS tether Mitofusin-2. This GTPase undergoes oxidation-dependent changes of function in mitochondrial dynamics, but whether this affects ERMCS is not clear (Shutt et al., 2012, Thaher et al., 2018).

To address this knowledge gap, we have focused on TMX2. This member of the TMX family has a cytosolic thioredoxin domain containing a SNDC motif (Guerra & Molinari, 2020). TMX2 mutations have been linked to a neurodevelopmental disorder with microcephaly, cortical malformations, and spasticity (NEDMCMS) (Ghosh et al., 2020, Vandervore et al., 2019). TMX2 can interact with ERMCS proteins like calnexin (Vandervore et al., 2019), but it is thought to lack a significant lumenal domain (Oguro & Imaoka, 2019). We therefore searched for interactors that could be targets on the cytosolic ERMCS face. Accordingly, we found that TMX2 modulates the redox state of mitochondrial TOM70 (translocase of the outer membrane 70). TOM70 not only promotes mitochondrial protein import, but can also tether the ER to mitochondria through interaction with IP_3_Rs (Filadi et al., 2018). Through its reductive activity, TMX2 prevents TOM70 sulfenylation and promotes redox-controlled ERMCS spacing, thereby interfering with ERMCS metabolic signaling. Our results describing the activity of TMX2 and its contributions for ERMCS functions provide important insight into NEDMCMS (Vandervore et al., 2019). Our fly *TMX2* knock-down animal model not only reproduces the NEDMCMS phenotype but also identifies glial cells as the point of origin of the pathology.

## Results

### TMX2 is a polytopic ERMCS transmembrane protein with a cytosolic SXXC motif

To address the mechanistic basis for NEDMCMS, we revisited the TMX2 topology and localization to determine where its putative catalytic domain might interact with substrates. From an AlphaFold three-dimensional structure model (Jumper et al., 2021), we determined that TMX2 likely exhibits four transmembrane domains across the ER membrane. These are preceded by an N-terminal membrane-apposed α-helix and followed by a cytosolic C-terminal thioredoxin domain (Figure 1A) containing a conserved SXXC_170_ motif (Figure 1B). As expected from the absence of potential glycosylation sites in the short lumenal domain, TMX2 is not glycosylated (Supplemental Figure 1A).

**Figure 1.**
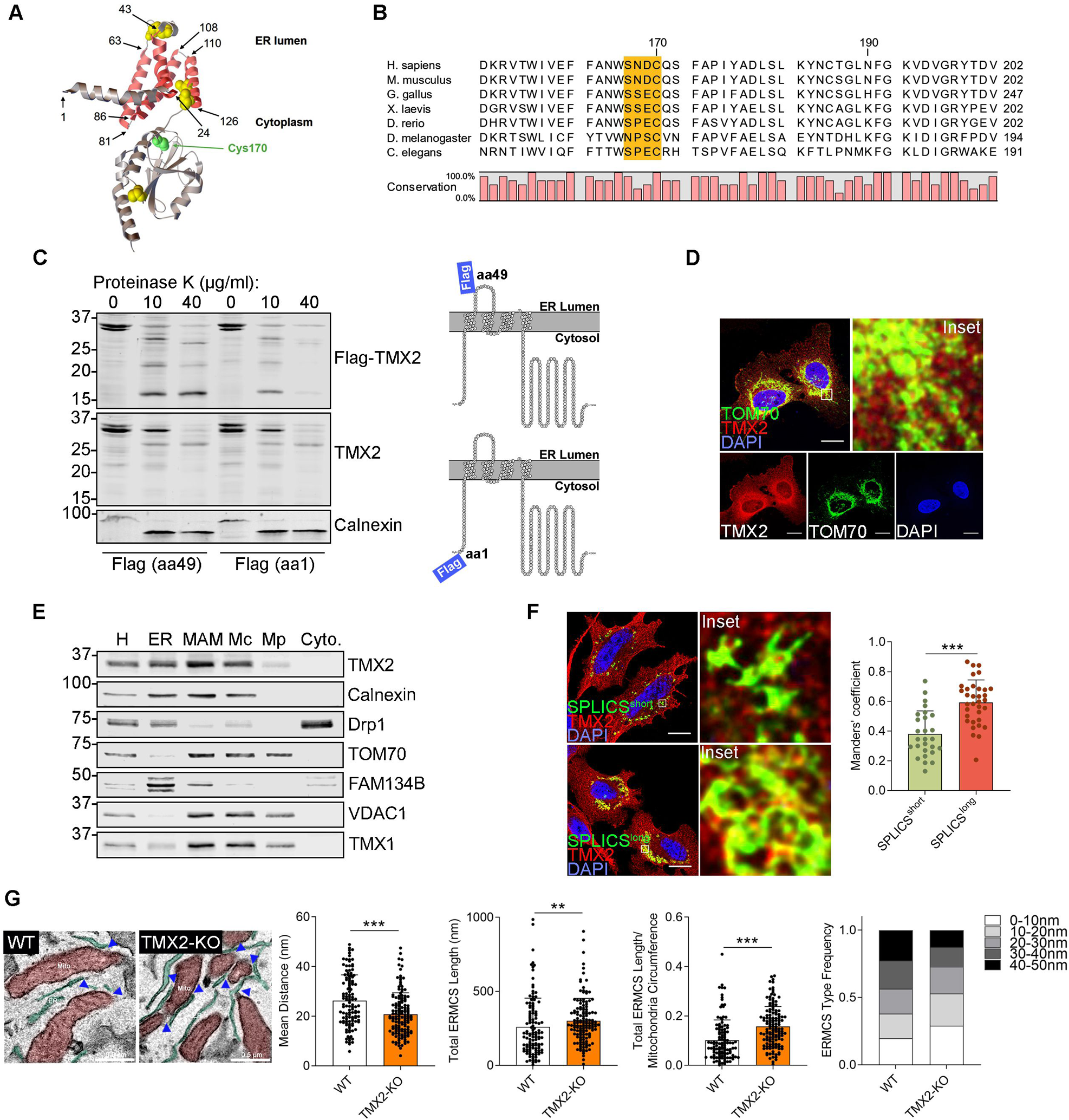
TMX2 is a ER-mitochondria contacts (ERMCS) protein that maintains contact site spacing. (A) Schematic overview of TMX2 membrane topology, as predicted by AlphaFold. Both N- and C-termini localize to the cytosol. Limits of transmembrane domains are indicated by numbers. Position of ER-lumenal and cytosolic portion are labeled. Cysteines are represented as yellow or green (catalytic C170) models. (B) Protein sequence alignment of the TMX2 thioredoxin domain portion containing the “SXXC” motif (highlighted in orange box) among species using the CLC Genomics Workbench (QIAGEN). (C) Proteinase K digestion of two TMX2 variants (see cartoon) with the Flag-tag inserted at amino acid 49 (version 1) or at the N-terminus (version 2). U251 MG cells expressing the two variants were incubated with different concentrations of proteinase K and then subjected to immunoblot analysis with Flag, TMX2, and calnexin antibodies. (D) Confocal immunofluorescence analysis of endogenous TMX2 in U251 MG cells. Overlap between TMX2 and TOM70 signal is indicated in yellow. Scale bar, 20 µm. (E) Percoll membrane fractionation of TMX2 of U251 MG cells. Protein components of subcellular fractions were analyzed by immunoblotting. Equal amounts were loaded and analyzed for the indicated MAM-regulatory proteins (H: homogenate; ER: endoplasmic reticulum; MAM: mitochondria-associated membrane; Mc: crude mitochondria; Mp: purified mitochondria; C: cytosol). (F) Co-localization with ERMCS SPLICS probes. HeLa cells were transfected with SPLICS-short (ERMCS distance 8-10 nm) or SPLICS-long (ERMCS distance 40-50 nm) plasmids and immunostained with TMX2 antibody. Mander’s coefficient analysis measured the co-localization between TMX2 and SPLICS signals (n = 27-34 of 2 technical replicates for each group, ***p < 0.001 by unpaired t-test). Scale bar, 20 µm. (G) Electron microscopy of TMX2 U251 MG wildtype and KO cells. Mitochondria were labeled in red, and ER was labeled in green. Average ER-mitochondria distance, and total ERMCS length were determined from electron micrographs (n = 75-91 of 3 technical replicates for each group, ***p < 0.001 by Mann-Whitney U test). ERMCS were classified and quantified by the mean distance. Scale bar, 0.5 µm.

Next, we investigated this postulated topology biochemically by proteinase K digestion and expressed TMX2 with a Flag-tag after the first transmembrane domain in the ER lumen (position 49; version 1) or a Flag-tag at the N-terminus (position 1; version 2) in HEK293 cells. Version 1 yielded a stable Flag-tagged 16 kDa fragment, corresponding to a Flag-tagged fragment with protected transmembrane and lumenal domains extending from proline 24 to methionine 125, with the short cytosolic domain from arginine 81 to histidine 86 (RSITV) likely embedded into the membrane (Figure 1C). In contrast, version 2 with the cytosolically exposed Flag tag, as well as the signal from an anti-TMX2 antibody directed towards the cytosolic domain were fully degraded (Figure 1C). Artificial glycosylation sites at residues 49 (L49N, projected lumenal), 71 (F71N, projected TM #2), N80 (R82T, projected TM #2), and N92 (F94S, projected TM #3, see Supplemental Figure 1B) all failed to generate glycosylated TMX2, in contrast to ERO1α used as a positive control (Supplemental Figure 1B). Therefore, TMX2 is a fully embedded four-transmembrane protein whose biggest extension corresponds to a cytosolic thioredoxin domain that is equipped with a classic KKXX ER retrieval motif at the C-terminus (Figure 1D).

Based on its co-fractionation with ERMCS markers VDAC1 and TMX1 on an Optiprep ER gradient (Lynes et al., 2012), we hypothesized that TMX2 is an ERMCS protein. Confirming this, we detected overlap between mitochondrial TOM70 and a validated anti-TMX2 antiserum in U251 MG cells (Figure 1D). A similar pattern was observed for the 3D reconstruction of co-localization between transfected Flag-tagged TMX2 and ERp57 or mitochondria in HeLa cells (Supplemental Figure 1C). Next, we quantitatively assayed TMX2 localization with a Percoll fractionation protocol that isolates mitochondria-associated membranes (MAMs) from the remainder of the ER. This showed that TMX2 is highly enriched in the MAM fraction, similar to calnexin, in both U251 MG (Figure 1E) and HeLa cells (Supplemental Figure 1D). ER and mitochondria association of TMX2 was not responsive to changes in redox conditions (Supplemental Figure 1E). To test whether TMX2 plays a role in ERMCS tethering, we next tested whether it is preferentially associated with tight (<10 nm) or widely spaced (>30 nm) ERMCS (Giacomello & Pellegrini, 2016). Thus, we quantified co-localization of endogenous TMX2 with SPLICS sensor proteins (Cieri et al., 2018), which specifically detect tight or wide ERMCS, respectively (SPLICS_short_ or SPLICS_long_, Figure 1F). This showed that TMX2 predominantly localizes to wide ERMCS. Next, we knocked out TMX2 in U251 MG cells by CRISPR/Cas9 (Supplemental Figure 1F). Transmission electron microscopy of these cells showed that TMX2 knockout increased the number of tight ERMCS, while it approximately halved the number of widely spaced ERMCS (Figure 1G). This was reproduced in U251 MG cells transfected with TMX2 RNAi (Supplemental Figure 1G, H). Together, our analyses showed that TMX2 provides a thioredoxin domain on the cytoplasmic face of widely spaced ERMCS.

### TMX2 Interacts with TOM70 to Decrease ERMCS Tethering

To further investigate the role of TMX2 for ERMCS, we used an unbiased mass-spectrometric approach to characterize the TMX2 interactome. Our crosslinked mass spectrometry (CLMS) showed that Flag-TMX2 interactors are found within a variety of functional clusters (Supplemental Figure 2A). Within this interactome, we were able to generate a protein-protein interaction network amongst multiple mitochondrial outer membrane (OMM) proteins that mediate ERMCS tethering and its regulation (Figure 2A), including members of the voltage-dependent anion channels (VDAC) (Reina et al., 2020, Shoshan-Barmatz et al., 2018), and TOM70 (Filadi et al., 2018).

**Figure 2.**
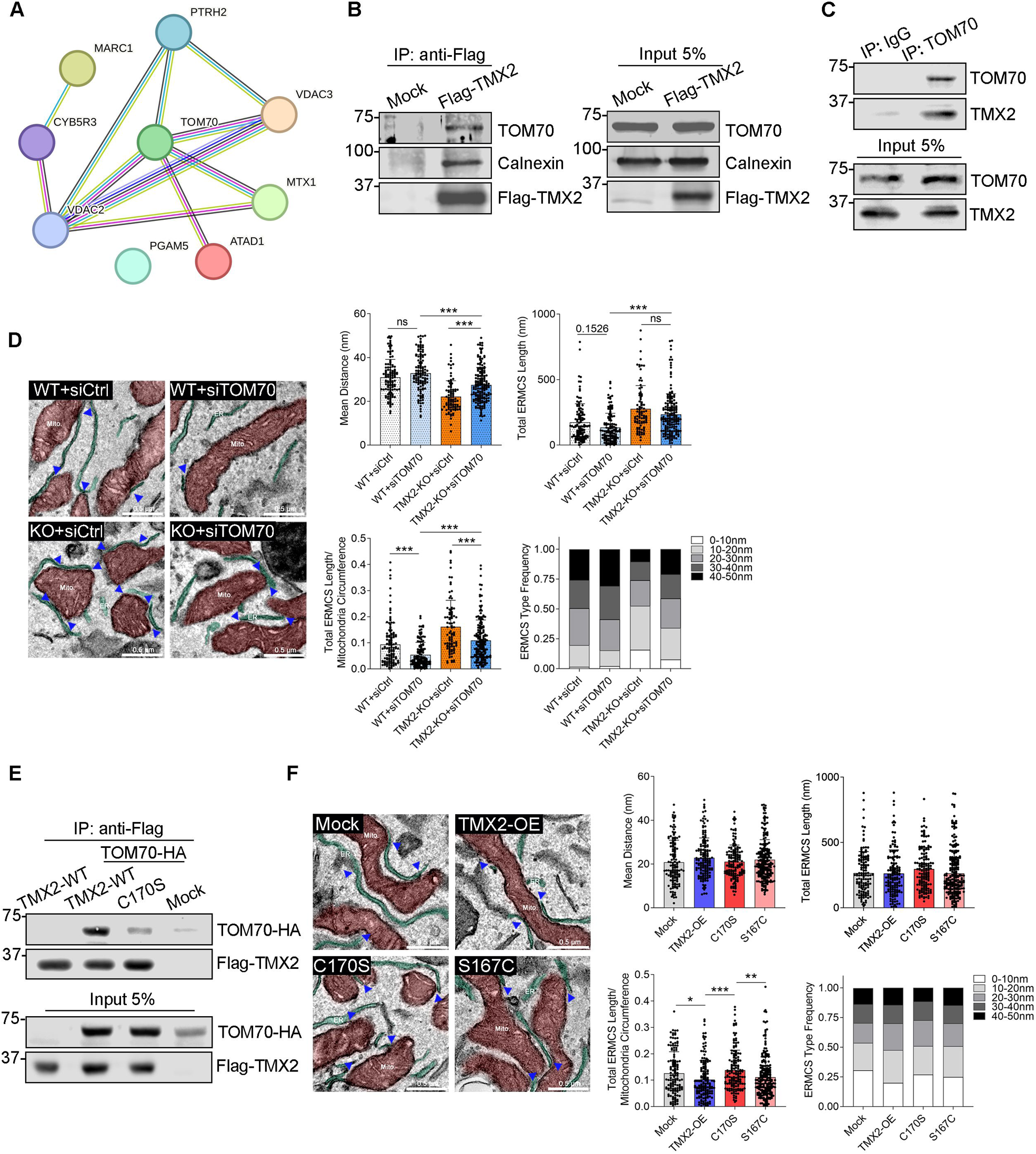
TMX2 interacts with TOM70 and promotes ERMCS spacing in a thioredoxin-like domain-dependent manner. (A) Mitochondrial outer membrane (MOM) interactome. MOM proteins were extracted from the interactome, and a protein-protein association network was generated with the STRING database (https://string-db.org). Line color indicates the evidence of a functional association. (B) Co-immunoprecipitation analysis. HEK293 cells were transfected with Flag-TMX2 for 48 h. The proteins co-immunoprecipitated with Flag-TMX2 were then analyzed for calnexin and TOM70. (C) Interaction between endogenous TMX2 and TOM70. Endogenous TOM70 was immunoprecipitated from HEK293 and analyzed for TMX2. (D) Transmission electron microscopy (TEM) analysis of TOM70 knock-down. wild-type and TMX2-KO U251 MG cells transfected with TOM70 RNAi (siTOM70) for 48 h. Mitochondria were labeled in red, and ER was labeled in green. Average ER-mitochondria distance, and total ERMCS length were quantified [n = 59-219 of 3 technical replicates for each group, *p < 0.05, **p < 0.01, and ***p < 0.001 by Kruskal-Wallis test (Dunn’s multiple comparisons test)]. The ERMCS were classified and quantified by the mean distance. Scale bar, 0.5 µm. (E) Co-immunoprecipitation analysis of wild-type and C170S TMX2. HEK293 cells were co-transfected with Flag-TMX2-WT, Flag-TMX2-C170S, and TOM70-HA for 24 h. The proteins co-immunoprecipitated with Flag-TMX2 were probed with an HA antibody. (F) Transmission electron microscopy (TEM) analysis of TMX over-expression. wild-type and TMX2-KO U251 MG cells transfected with TMX2 constructs as indicated for 48 h. Mitochondria were labeled in red, and ER was labeled in green. Average ER-mitochondria distance, and total ERMCS length were quantified [n = 59-219 of 3 technical replicates for each group, *p < 0.05, **p < 0.01, and ***p < 0.001 by Kruskal-Wallis test (Dunn’s multiple comparisons test)]. The ERMCS were classified and quantified by the mean distance. Scale bar, 0.5 µm.

TOM70 is enriched on ERMCS, where it acts to promote tethering (Filadi et al., 2018) and is known to be redox-responsive in yeast (Kreimendahl et al., 2020). This made it a good candidate target for TMX2. We validated their interaction through co-immunoprecipitation of endogenous TOM70 with Flag-tagged TMX2 (in parallel to the known TMX2/calnexin interaction, Figure 2B) and with endogenous TMX2 (Figure 2C). Moreover, it has spatial consequences. While reduction of TMX2 levels increases ERMCS (<10 nm, Figure 1G), ERMCS partially revert to normal upon depletion of both TMX2 and TOM70 (Figure 2D, Supplemental Figure 2B).

Next, we investigated the requirements of the TOM70-TMX2 interaction. This analysis showed that mutation of C170, the sole cysteine within the TMX2 active site, abrogated co-immunoprecipitation (Figure 2E). This dependence on C170 was also observed for calnexin but not for VDAC1 (Supplemental Figure 2C). We further investigated whether ERMCS spacing depends on TMX2 by showing that plasmid-based over-expression of wildtype TMX2 (Supplemental Figure 2D) increased ERMCS spacing, manifesting as a more than 30% reduction of tight ERMCS upon over-expression (Figure 2F). Importantly, this change did not occur with the C170S mutant, thus demonstrating that a cytosolic thioredoxin domain is instrumental in the ability of TMX2 to increase ERMCS spacing in a TOM70-dependent manner.

### TMX2 Prevents TOM70 Sulfenylation with the Help of PRDX1

To understand the consequence of TMX2 for ERMCS, we characterized the TMX2-TOM70 interaction and its function. Given the large size of the cytosolic domain of TOM70, we constructed deletion mutants of the chaperone-binding (residues 138-251) and substrate-binding (residues 468-608) domains. Deletion proteins of the core domain (residues 292-468) were poorly expressed and showed drastically altered mitochondrial structure (Supplemental Figure 3A). In contrast, deletion mutants of the chaperone-binding domain were well expressed but did not allow for TMX2 interaction (Supplemental Figure 3B). Interestingly, the cysteine-rich chaperone-binding domain was previously identified as critical to prevent mitochondrial stress in yeast (Backes et al., 2021), suggesting TMX2 affects the functions of TOM70 in ERMCS tethering or mitochondrial protein import.

Unlike their typically oxidizing ER-lumenal counterparts, cytosolic thioredoxin proteins can directly prevent protein oxidation (Arner & Holmgren, 2000) or promote PRDX antioxidant functions (Netto & Antunes, 2016). To investigate whether TMX2 fits into the ER lumenal or cytosolic thioredoxin type, we first used a redox-sensitive GFP (ro-GFP2)-based assay as a qualitative readout for TMX2 activity. Genetic fusion constructs between roGFP2 and thioredoxin superfamily proteins have previously been used to study structure-function relationships in both PRDXs and glutaredoxins (Liedgens et al., 2020, Staudacher et al., 2018, Zimmermann et al., 2021). Expressing a roGFP2-TMX2_cytosolic_ construct in specifically engineered yeast cells, TMX2 neither showed a thiol peroxidase-like nor a glutaredoxin-like activity, in contrast to human glutaredoxin-1 (Supplemental Figure 3C).

To determine whether the TMX2-TOM70 interaction could instead prevent oxidation of TOM70, we turned to the DCP-Bio1 compound, a reagent that specifically detects sulfenylated cysteines (Poole et al., 2007) (Figure 3A). We detected threefold increased DCP-Bio1-labeled TOM70 signals in siTMX2-transfected HEK293 cells (Figure 3B). This was confirmed by mass-spectrometric analysis that detected C206 sulfenylation (Figure 3C). The oxidation of TOM70 in TMX2 knock-down cells required ROS production in mitochondria and the ER from OXPHOS complex III and NOX4 (Supplemental Figure 3D). To distinguish between a passive oxidation or oxidation involving PRDXs, we tested whether TMX2 can interact with cytosolic PRDX. Indeed, through CLMS, we could detect PRDXs 1, 2 and 6 within the TMX2 interactome (Figure 3D). This interaction was confirmed by co-immunoprecipitation for PRDX1 (Figure 3E).

**Figure 3.**
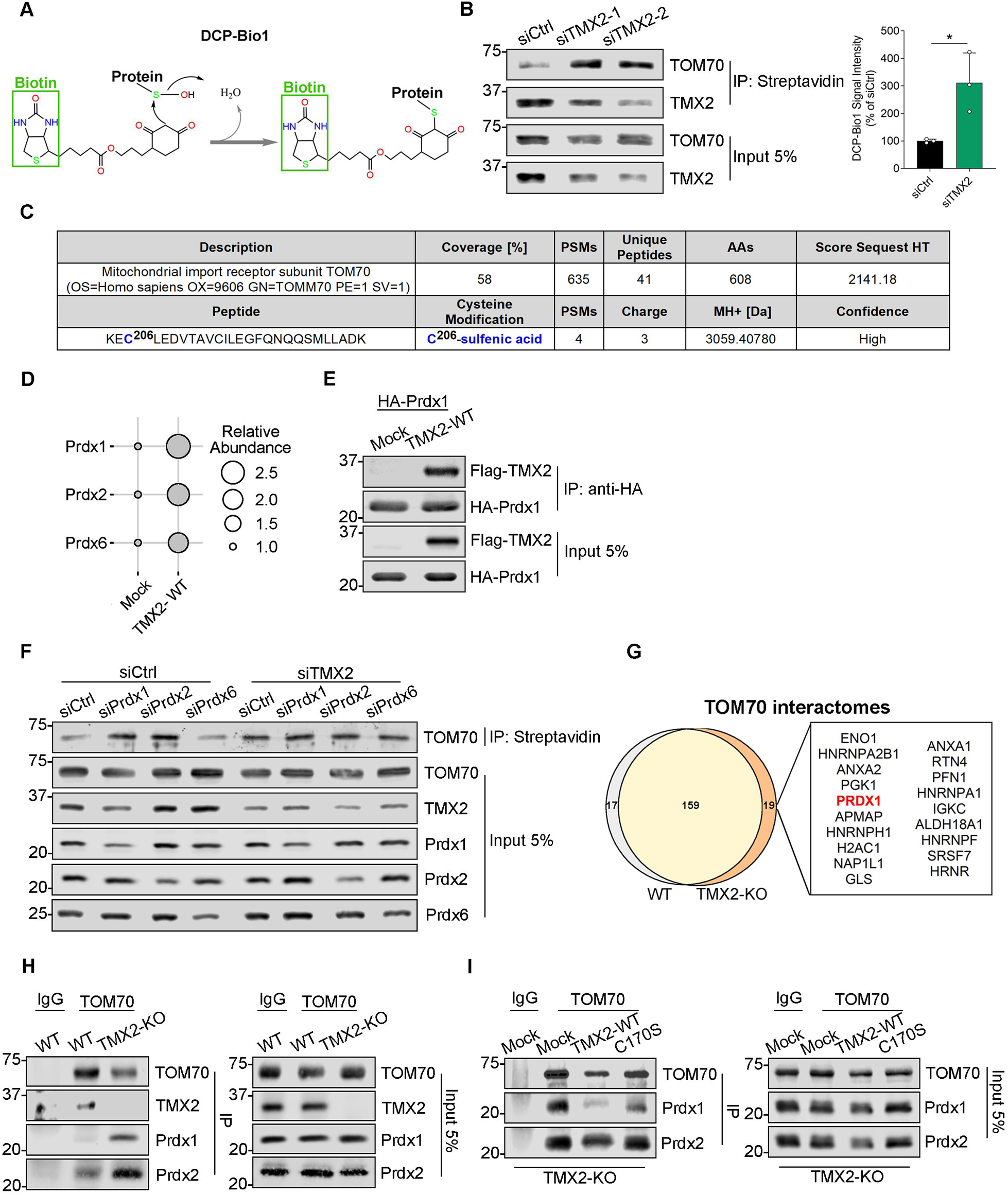
TMX2 prevents TOM70 sulfenylation through PRDX1. (A) Cartoon depicting labeling of sulfenylated proteins with DCP-Bio1. The DCP-Bio1 reactive carbon covalently attaches to a protein at its sulfenylation site. (B) DCP-Bio1 measurement of TOM70 sulfenylation in the presence of TMX2 RNAi. The assay was performed in HEK293 cells after transfecting with TMX2 RNAi for 48 h. The sulfenylated cysteine residues were biotinylated and immunoprecipitated with streptavidin beads. The immunoprecipitated proteins were then analyzed by Western blotting with the indicated antibodies [n = 3 biological replicates for each group, *p < 0.001 by unpaired t-test]. (C) Mass spectrometric analysis of sulfenylation modification within immunoprecipitated TOM70. HEK293 isolates from control and TMX2 RNAi cells were used to detect redox modifications, C206 was identified (n = 3 biological replicates). (D) PRDX interactions within the TMX2 proteome. Interacting PRDXs were detected via CO-IP-MS analysis and visualized as a bubble plot. The size of each bubble represents the relative abundance normalized to the empty vector group. (E) PRDX1 interaction with TMX2. HEK293 cells were co-transfected with Flag-TMX2 and HA-PRDX1 for 24 h. The proteins co-immunoprecipitated with HA-PRDX1 were probed with a Flag antibody. (F) DCP-Bio1 measurement of TOM70 sulfenylation in the presence of TMX2 and PRDX RNAi. HEK293 cells were transfected with the indicated RNAi for 48 h. Then the DCP-Bio1 assay was performed and the immunoprecipitated proteins were analyzed by Western blots. (G) TOM70 interactome in wild-type and TMX2 U251 MG KO cells. CO-IP-MS analysis was subjected to background subtraction, and the interactomes were plotted in a Venn diagram. (H) PRDX1 interaction with TOM70 dependent on TMX2. In wild-type and TMX2-KO cells, the proteins co-immunoprecipitated with the endogenous TOM70 were analyzed by Western blotting with the indicated antibodies. (I) PRDX1 interaction with TOM70 dependent on TMX2 C170. TMX2-KO cells were transfected with Flag-TMX2 and C170S plasmids for 48 h. Then the co-immunoprecipitation was performed as described above.

Next, we tested whether PRDXs functionally interact with TMX2 to prevent TOM70 sulfenylation. Consistent with such an activity, we found TOM70 sulfenylation increased upon PRDX1 and PRDX2, but not PRDX6 knock-down (Figure 3F). If TMX2 uses PRDX1 and PRDX2 to prevent TOM70 sulfenylation, their binding to TOM70 should increase in the absence of TMX2. We therefore screened the TOM70 interactome of TMX2 wild-type and KO U251 MG cells for interacting PRDXs and detected PRDX1 bound to TOM70 in KO cells (Figure 3G). Increased interaction between TOM70 and PRDX1 was also detected in KO over wild-type cells by co-immunoprecipitation (Figure 3H). This interaction was prevented by the transfection of wild-type TMX2 but not by C170S TMX2 (Figure 3I). For PRDX2, we could only detect minor changes in the interaction pattern. Therefore, TMX2 promotes the PRDX1 and PRDX2 antioxidant activities to prevent TOM70 sulfenylation from mitochondrial and ER ROS.

### TMX2 Decreases the Mitochondrial Ca^2+^ Response

We hypothesized that TMX2-controlled ERMCS spacing impacts this membrane contact not just spatially but also functionally. Therefore, we next analyzed the protein content of ERMCS in U251 MG TMX2 KO cells. We took advantage of a recently published, detailed description of the ERMCS proteome that is based on a split BioID approach (Kwak et al., 2020) and quantified changes of *bona fide* ERMCS proteins within ERMCS isolates from U251 MG TMX2 wild-type and KO cells through LC-MS analysis. We found that most ERMCS proteins (e.g., PDIA6, VAPB, TMX1) increased significantly except for FUNDC2 (Figure 4A) and that, overall, the number of proteins associated with ER protein folding and ER membrane organization increased in parallel (Supplemental Figure 4A), a phenotype known to promote ER-mitochondria Ca^2+^ flux in multiple examples (Simmen & Herrera-Cruz, 2018).

**Figure 4.**
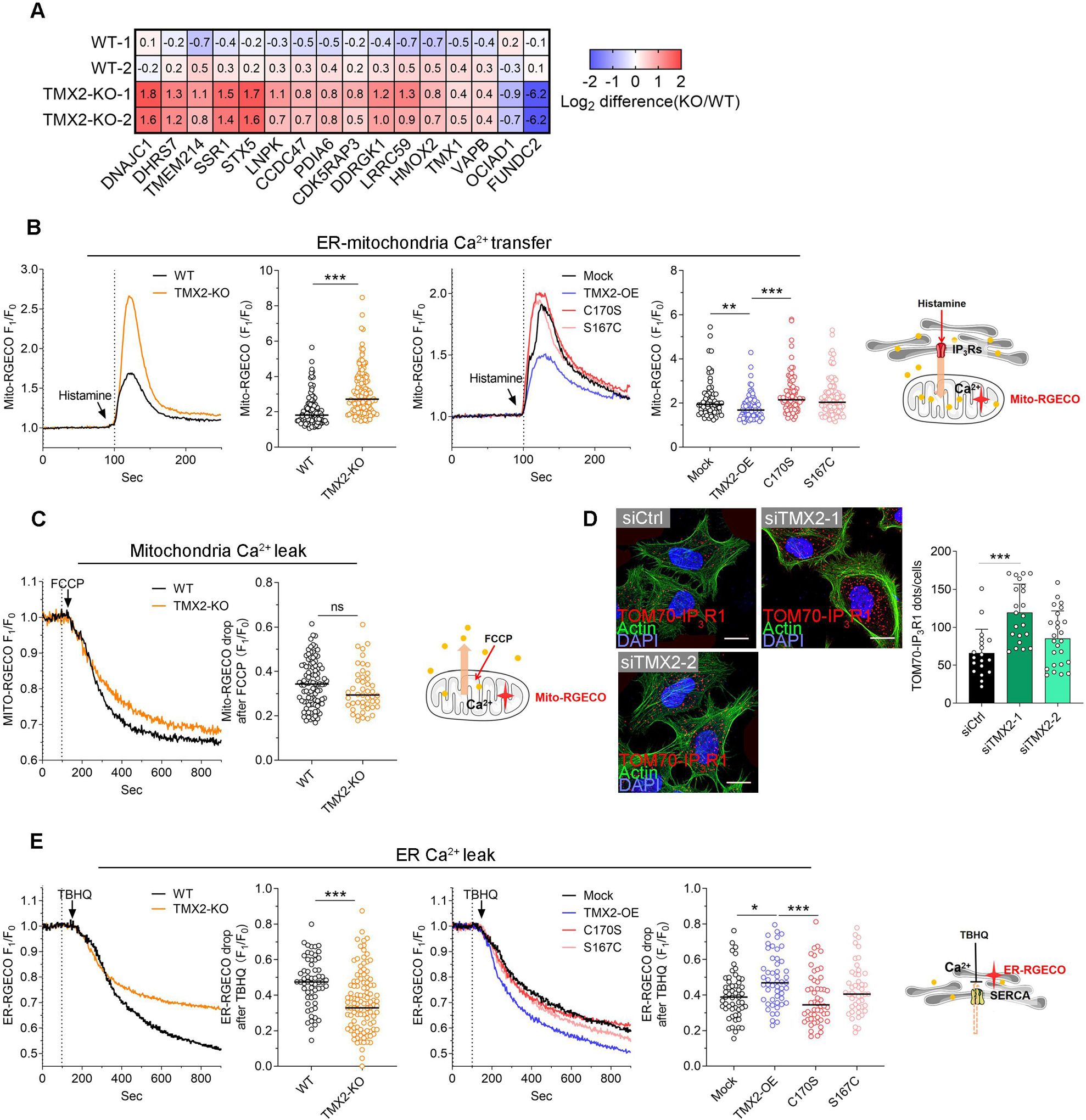
TMX2 alters ERMCS and limits ER-mitochondria Ca^2+^ transfer. (A) MAM proteome of U251 MG TMX2 wild-type and KO cells. MAM fractions were isolated from wild-type and TMX2-KO cells by Percoll gradient fractionation, followed by LC-MS analysis to detect the MAM proteomics altered under TMX2 depletion. The samples were then loaded on a precast gel (50 µg protein each), digested, and the tryptic peptides were identified by Proteome Discoverer 2.5. Data was normalized to each sample’s total reconstructed peak area and the abundance of identified proteins was represented by a heatmap (n = 2 biological replicates). (B) ER-mitochondria Ca^2+^ transfer measurements. Wild-type and TMX2-KO cells and U251 cells stably transfected with TMX2 and its mutants were transfected with the mitochondria-targeted and Ca^2+^ sensitive fluorescent probe Mito-RGECO for 24 h. The Ca^2+^ signal was induced with 50 µm histamine (IP_3_R activator). Relative fluorescence increases were recorded and quantified [n = 62-143 of 3-4 biological replicates for each group, **p < 0.01, and ***p < 0.001 by Mann-Whitney U test or Kruskal-Wallis test (Dunn’s multiple comparisons test)]. (C) Mitochondria Ca^2+^ content. Wild-type and TMX2-KO cells were transfected with Mito-RGECO for 24 h, followed by incubating with 10 µM FCCP to block mitochondrial Ca^2+^ import. Live-cell images were recorded and quantified (n = 47-96 of 3-4 biological replicates for each group, *p < 0.05 by Mann-Whitney U test). (D) Representative immunofluorescence images of the proximity ligation assay (PLA) between IP_3_R1 and TOM70. HeLa cells were transfected with TMX2 RNAi for 48 h and subjected to the assay. The protein interactions (within 40 nm), visualized as a single dot, were counted [n = 19-25 of 2 technical replicates for each group, ***p < 0.001 by one-way ANOVA (Dunnett’s multiple comparisons test)]. Scale bar, 20 µm. (E) ER Ca^2+^ content. ER Ca^2+^ leak under TBHQ (SERCA pumps inhibitor) treatment was performed in wild-type and TMX2-KO cells, and U251 cells stably transfected with TMX2 and its mutants after transfecting with ER-RGECO for 24 h. The drop of the fluorescent signal is proportional to the ER Ca^2+^ content. The fluorescence was recorded and quantified [n = 47-122 of 3-4 biological replicates for each group, *p < 0.05, and ***p < 0.001 by Mann-Whitney U test or Kruskal-Wallis test (Dunn’s multiple comparisons test)].

Thus, we tested whether this was indeed the case. Our results showed that histamine-triggered Ca^2+^ transfer from the ER to mitochondria increased by about 50% in TMX2 KO (Figure 4B) or knock-down cells (Supplemental Figure 4B). In contrast, over-expression of wildtype TMX2 but not of the C170S mutant decreased Ca^2+^ transfer (Figure 4B). Since we detected changes in Ca^2+^ flux, we next released mitochondrial Ca^2+^ with FCCP that dissipates the mitochondrial membrane potential and blocks store-operated Ca^2+^ entry (Yoast et al., 2021). This was not affected by TMX2 (Figure 4C, Supplemental Figure 4C). Consistent with TMX2-controlled Ca^2+^ communication, a proximity ligation assay (PLA) for the TOM70-IP_3_R1 interaction pair detected increased signals upon TMX2 knock-down in HeLa cells (Figure 4D). These results suggested that TMX2 prevents excessive Ca^2+^ transfer between the ER and mitochondria but does not significantly alter mitochondrial Ca^2+^ content.

Such an effect could be based on increased ER Ca^2+^ content or facilitated Ca^2+^ release by TMX2, as reported (Gutierrez et al., 2020, Raturi et al., 2016). While reduced levels of TMX2 did not alter ER Ca^2+^ release dynamics (Supplementary Figure 4D), they reduced ER Ca^2+^ content (Figure 4E, Supplementary Figure 4E). In contrast, TMX2 over-expression increased ER Ca^2+^ content, dependent on the presence of C170 within its thioredoxin domain (Figure 4E). Therefore, TMX2 increases ER Ca^2+^ content and inhibits ER-mitochondria Ca^2+^ transfer.

### TMX2 Maintains the Mitochondrial Lipidome and Reduces ROS from OXPHOS

ERMCS do not only determine Ca^2+^ signaling, they also control the global cellular lipidome such as phosphatidylcholine (PC) levels (Ganji et al., 2023). For our analysis, we decided to focus on lipid species with known functional connections to ERMCS (Vance, 2020) and analyzed TMX2 U251 MG wildtype and KO cells through LC-MS and LC-MS/MS. In addition to PC, we included phosphatidylserine (PS) that is converted to phosphatidylethanolamine (PE) within mitochondria (Vance, 2015) and phosphatidic acid (PA), which determines the mitochondrial cardiolipin (CL) content (Yeo et al., 2021), all of which are under the control of ERMCS (Perea et al., 2023). This showed that most changes occurred amongst the PC and Hexosylceramide (HexCer) species (Supplemental Figure 5A). When inspecting a lipidome volcano plot, we found that several CL species increased in KO cells (Figure 5A). A more detailed analysis showed that PC and PE species of the ether subtype were decreased (Figure 5B, C), which is known to increase mitochondrial ROS production (Chen et al., 2023).

**Figure 5.**
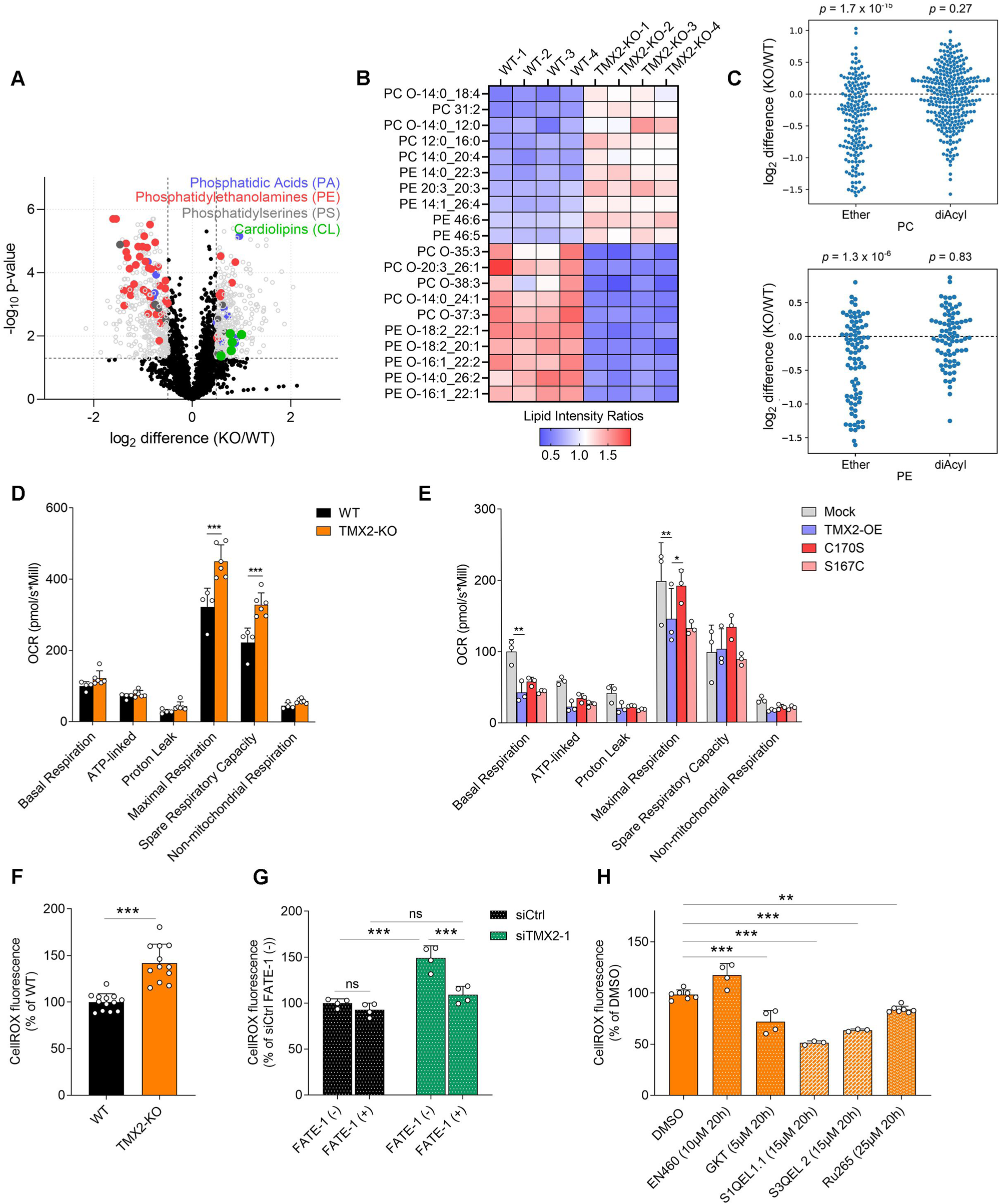
TMX2 regulates cellular lipidomics and mitochondrial respiration. (A) U251 MG TMX2 KO lipidomics. Altered lipid species in U251 MG TMX2-KO cells (versus wild-type) were visualized on a volcano plot. For each lipid molecule species, the abundance change between wild-type and TMX2-KO cells on the x-axis against the p-value on the y-axis was plotted. The significance was determined as FC >1.40 or <0.71, p-value <0.05, and p-value adjusted for false-discovery rate (q-value) <0.25. The analysis showed 347 significantly up-regulated lipids with FC >1.40 and 370 significantly down-regulated lipids with FC <0.71 (n = 4 biological replicates for each group, analyzed by unpaired t-test with Welch correction). (B) Heatmap for the relative abundance of the top 5 significantly altered PC and PE species. (C) Individual lipid species changes for PC (top) and PE (bottom). PC and PE subtypes were classified (n = 4 biological replicates for each group, analyzed by one-sample t-test). (D, E) Analysis of oxygen consumption rates in wild-type and TMX2-KO cells and U251 MG cells stably expressing TMX2 mutants [n = 3 biological replicates for each group, *p < 0.05, **p < 0.01, and ***p < 0.001 by two-way ANOVA (Dunnett’s multiple comparisons test)]. (F) Cytosolic ROS production in wild-type and TMX2-KO cells. CellROX™ Orange values were plotted (n = 3-4 biological replicates for each group, ***p < 0.001 by unpaired t-test). (G) Cytosolic ROS production dependent upon the ERMCS spacer FATE-1. U251 MG cells were transfected with indicated siRNA constructs for 24 h and further transfected with FATE-1-GFP for 48 h, and CellROX™ Orange values were plotted [n =3 biological replicates for each group, ***p < 0.001 by two-way ANOVA (Šídák’s multiple comparisons test)]. (H) Cytosolic ROS production upon ER and mitochondrial ROS inhibition. TMX2-KO cells were treated with 10 µm EN460 (ERO1 inhibitor), 5 µm GKT (NOX4 inhibitor), 15 µm S1QEL1.1 (Complex I inhibitor), 15 µm S3QEL 2 (Complex III inhibitor), and 25 µm Ru265 (MCU inhibitor) overnight before analysis [n = 2-3 biological replicates for each group, **p < 0.01, and ***p < 0.001 by one-way ANOVA (Dunnett’s multiple comparisons test)] CellROX™ Orange values were plotted.

Together, the promoted Ca^2+^ signaling (Cardenas et al., 2010) and the increase of CL (Zhang et al., 2002) predicted that TMX2 KO cells should show increased mitochondrial respiration but at the cost of increased ROS production. Accordingly, we found that TMX2 depletion increased maximal respiration by about 30% (Figure 5D, Supplemental Figure 5B). The inhibition of mitochondrial Ca^2+^ import with Ru265 reversed this increase in full, while TOM70 knock-down partially reduced it (Supplemental Figure 5C). Conversely, TMX2 over-expression reduced maximal respiration by about 20% compared to control cells (Figure 5E). These changes in respiration were accompanied by increased ROS production upon TMX2 depletion (Figure 5F, Supplemental Figure 5D). Conversely, ROS decreased in TMX2 over-expressing cells, but not with C170S (Supplemental Figure 5E). Mitochondrial dysfunction could derive from abnormal ERMCS spacing, or from altered TOM70 oxidation. To distinguish between these possibilities, we first introduced FATE-1 to increase ERMCS spacing. This abolished increased ROS production completely (Figure 5G). Next, we depleted TOM70 in wild-type and TMX2 KO cells. This experiment showed that TOM70 knock-down overall decreased ROS but did not result in a difference between TMX2 wild-type and KO U251 MG cells (Supplemental Figure 5F). Importantly, the ROS observed from TMX2 KO were derived from ER (NOX4) and mitochondrial sources (complex I and III, Figure 5H). These findings indicate that the mitochondrial dysfunction, characterized by altered respiration and increased ROS production, is connected to ERMCS spacing and proper mitochondrial Ca^2+^ homeostasis, functions partially dependent on TOM70.

### TMX2 Preserves the Mitochondrial Proteome

To test whether TMX2 preserves normal mitochondrial functions through maintenance of the mitochondrial proteome, we biochemically isolated the mitochondria proteome from U251 MG wildtype and TMX2 KO cells and analyzed it by mass spectrometry (Figure 6A). This revealed that several mitochondrial proteins decreased in TMX2 KO cells. A subsequent treatment with proteinase K to eliminate surface proteins revealed significant decreases in OXPHOS (e.g., NDUFB9, and COX5B) and Krebs cycle components (e.g., SUCLG1). These decreases were not uniform (Supplemental Figure 6A), but the decrease in mitochondrial protein content was most significant for proteins of the mitochondrial inner membrane (MIM, Figure 6B). Within the OXPHOS machinery, components of OXPHOS complex IV and V decreased, while OXPHOS complex II showed an increase (Figure 6C). We verified this pattern in total lysates for select mitochondrial proteins and found the expected changes for representative complex I-V proteins (Figure 6D). These changes observed within KO cells could not be reversed by increased ERMCS spacing through FATE-1 (Supplemental Figure 6B). An obvious explanation for these findings could be the known function of TOM70 in mitochondrial protein import. To investigate this question, we took two approaches: 1) we tested whether known TOM70 importome proteins (Morgenstern et al., 2021) show consistently decreased interaction with TOM70 in the absence of TMX2. This showed that some of these proteins like the mitochondrial fission and fusion controlling ARMC10 (Chen et al., 2019) indeed interacted less with TOM70 in TMX2 KO cells. However, many others (e.g., DLAT) did not (Figure 6E). 2) We tested whether the transcription of some of the most changed proteins was altered in TMX2 KO cells. This also did not reveal a trend (Figure 6F), suggesting that TMX2 modulates the mitochondrial proteome independent of TOM70.

**Figure 6.**
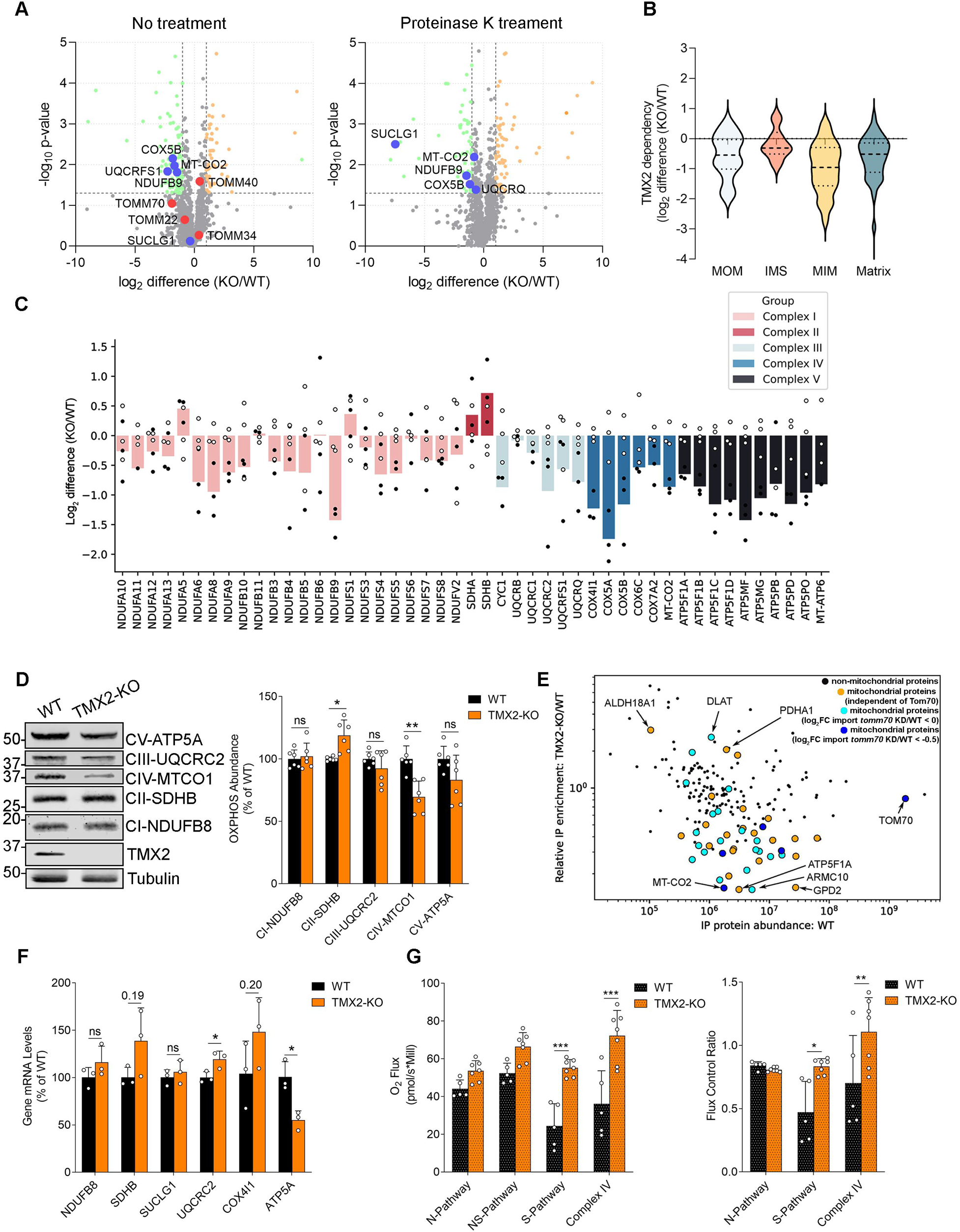
TMX2 maintains the normal mitochondrial proteome and function. (A) Changes in the mitochondrial proteome in U251 MG TMX2 KO cells. Volcano plots of mitochondrial proteomics difference between wild-type and TMX2-KO cells. MitoCarta3.0 was used to filter mitochondrial proteins, and representative TOM70 substrates. The significance was determined as FC >2 or <0.5, and p-value <0.05 (n = 3 biological replicates for each group, analyzed by unpaired t-test with Welch correction). (B) Further classification of mitochondrial proteome changes. Plots from A from wild-type and TMX2-KO cells were further classified into subgroups (MOM: Outer mitochondrial membrane; IMS: mitochondrial intermembrane space; MIM, Inner mitochondrial membrane; and mitochondrial matrix). (C) Further classification of OXPHOS protein changes. The electron transport chain (ETC) components were extracted from the dataset and analyzed (WT: open circles; TMX2-KO: filled circles). (D) Western blot analysis of OXPHOS protein changes. The cell lysates from wild-type and TMX2 were collected and analyzed with Western blots as indicated (n= 3 biological replicates for each group, *p < 0.05, and **p < 0.01 by multiple unpaired t-test). (E) Correlation of TOM70 protein interactions in U251 MG wild-type and TMX2 KO cells with TOM70 importome. Within the interactome, TOM70 interacting mitochondrial proteins were determined (MitoCarta 3.0) were determined in wild-type and TMX2-KO cells by CO-IP-MS analysis. The interactome was correlated to the TOM70 importome (see text) [Black: non-mitochondrial proteins; Orange: localized to mitochondria independently of TOM70; Light blue: localized to mitochondria, dependent on TOM70 (log_2_FC import *TOM70* KD/wild-type < 0); Dark blue: localized to mitochondria, dependent on TOM70 (log_2_FC import *TOM70* KD/wild-type < −0.5)]. (F) Transcription of select OXPHOS proteins. Gene expression alterations of OXPHOS proteins between U251 MG wild-type and TMX2 KO cells were tested by RT-qPCR analysis (n= 3 biological replicates for each group, statistics were performed via multiple unpaired t-test). (G) OXPHOS pathway analysis. Wild-type and TMX2-KO cells were permeabilized with digitonin, and then the mitochondrial respiration assays were performed (see Figure S6F). The contribution of each ETC pathway (N, NADH; NS, NADH-Succinate; S, Succinate) to O_2_ flux and flux control ratio was measured and plotted [n = 3-4 biological replicates for each group, *p < 0.05, **p < 0.01, and ***p < 0.001 by two-way ANOVA (Šídák’s multiple comparisons test)].

To further examine the bioenergetic role of TMX2, we measured the mitochondrial membrane potential and found it was lower upon TMX2 depletion (Supplemental Figure 6C). In addition, we detected that AMP-activated protein kinase (AMPK) phosphorylation levels were unchanged in TMX2 KO cells (Supplemental Figure 6D), correlating with the inability of TMX2 KO cells to rescue their low ATP levels (Supplemental Figure 6E). Within mitochondria of TMX2 KO cells and to a lesser extent upon TMX2 RNAi transfection, O_2_ flux increased most within complex IV and the succinate (S)-pathway (Gnaiger, 2009) (Figure 6G, Supplemental Figure 6F, G). The latter observation correlates with the observed specific increase of complex II proteins (Figure 6C), which accept succinate from the Krebs cycle (Lemieux et al., 2017).

### TMX2 Patient Mutations and *in vivo* Knock-down Compromise TMX2 Function

Previous reports on TMX2 have suggested that the autosomal-recessive NEDMCMS pathology derives from a loss of function. This is based on *TMX2* truncating variants observed in patients and the reduction of TMX2 protein levels in patient samples (Vandervore et al., 2019). The present study provides the first comprehensive, functional analysis of TMX2. We sought to investigate some of the key functions we have herein identified in *TMX2* variant cells from NEDMCMS patients and *TMX2* knock-down in an animal model to determine the pathogenesis of NEDMCMS. We initiated our characterization with three NEDMCMS patient fibroblasts. These cell lines were labeled P1 (male, transheterozygous D55A/²1136-296); P4 (male, homozygous G56A), and P5 (female, homozygous G56A) (Vandervore et al., 2019) (Supplemental Figure 7A). TMX2 protein levels were reduced in P1, while they were comparable in P4 and P5 (Figure 7A). We confirmed that the mitochondrial proteins under the control of TMX2 were decreased (e.g., TOM70, SUCLG1). AMPK phosphorylation was increased in P4 and P5, but not P1 (Figure 7A). Therefore, NEDMCMS patient cells partially recapitulate TMX2 depletion results (Figure 6). Regarding the mitochondrial proteins of complexes I-V, this mirroring was also incomplete and was restricted to decreases in complex III UQCRC2 in P4 and P5 (Supplemental Figure 7B). We proceeded with a functional characterization of the defects in NEDMCMS patient cells and detected increased ERMCS in P1 but not P4 and P5 (Figure 7B), increases in maximal respiration (Figure 7C), and increases of various extent in cellular ROS (Figure 7D). Unlike the decrease in mitochondrial membrane potential observed upon TMX2 depletion (Supplemental Figure 6E), we detected an increase in P1 and P5, while P4 showed no change (Supplemental Figure 7C), accompanied by an insignificant decrease in total ATP (Supplemental Figure 7D). Therefore, NEDMCMS patient cells do not provide an unequivocal loss of function phenotype but are rather characterized by complex ERMCS dysfunction, where respiration and ROS matched the TMX2 depletion phenotype.

**Figure 7.**
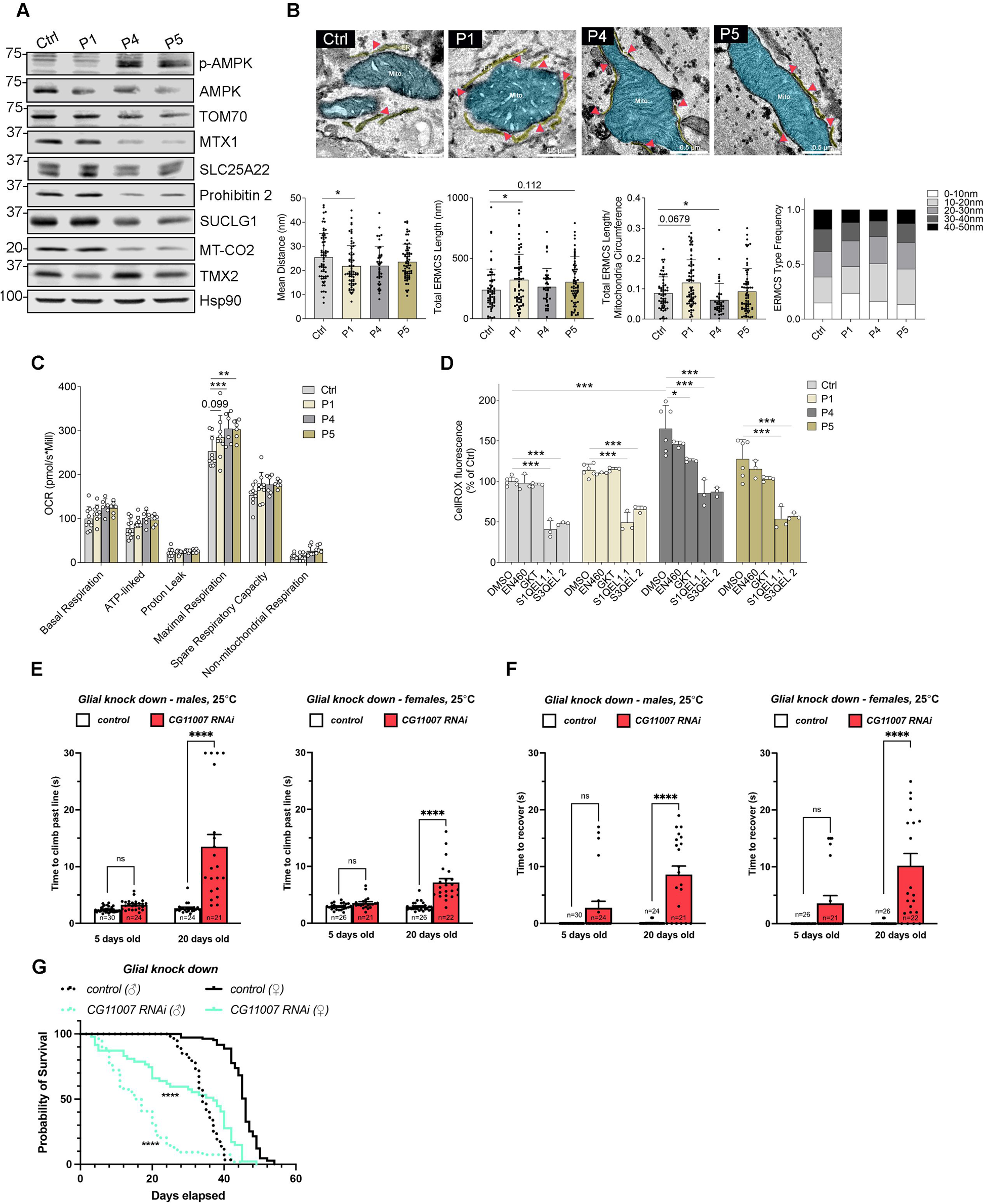
*TMX2* mutant patient fibroblasts and glial *TMX2* knock-down *D. alanogaster* animals exhibit partial TMX2 loss of function. (A) Lysate analysis from control and NEDMCMS patient fibroblasts. Cell lysates were collected and analyzed by Western blots with the indicated antibodies. (B) Representative electron microscopy images of NEDMCMS patient fibroblasts. Mitochondria were labeled in blue, and ER was labeled in yellow. Mitochondria circumference, average ER-mitochondria distance, and total ERMCS length were measured from electron micrographs [n = 27-71 of 3 technical replicates for each group, *p < 0.05, and ***p < 0.001 by Kruskal-Wallis test (Dunn’s multiple comparisons test)]. The ERMCS were classified and quantified by the mean distance. Scale bar, 0.5 µm. (C) Oxygen consumption rates in NEDMCMS patient fibroblasts. High-resolution respirometry was performed to measure mitochondrial respiration of NEDMCMS patient fibroblasts [n = 3-4 biological replicates for each group, **p < 0.01, and ***p < 0.001 by two-way ANOVA (Šídák’s multiple comparisons test)]. (D) Cytosolic ROS levels in NEDMCMS patient fibroblasts. Patient fibroblasts were treated with 10 µm EN460 (ERO1 inhibitor), 5 µm GKT (NOX4 inhibitor), 15 µm S1QEL1.1 (Complex I inhibitor), 15 µm S3QEL 2 (Complex III inhibitor), and 25 µm Ru265 (MCU inhibitor) overnight. Then the cytosolic ROS production was measured by flow cytometry [n = 3-6 biological replicates for each group, *p < 0.05, and ***p < 0.001 by two-way ANOVA]. CellROX™ Orange values were plotted. (E, F) Pan-glial knock-down of *CG11007* (repo-GAL4 > UAS-CG11007-RNAi) at 25°C causes age-dependent climbing deficits and mechanically-induced seizures in male and female flies at 20 days post-eclosion. This phenotype is not observed in control flies (repo-GAL4 > UAS-luciferase-RNAi) or young animals at 5 days post-eclosion [n = indicated in figure for each group, ****p < 0.0001 by one-way ANOVA (Tukey post-hoc test)]. (G) Life span analysis of flies with pan-glial knock-down of *CG11007*. Kaplan Meier graph of male and female animals, as indicated. ****p < 0.0001 by one-way ANOVA (Tukey post-hoc test).

Next, we investigated which cell type could compromise nervous system functions in patients. To approach this question, we took advantage of the high degree of conservation observed for TMX2 (Figure 1A, B) and chose the *D. melanogaster* fly model. The uncharacterized *Drosophila* gene, *CG11007*, is the single ortholog to human *TMX2* having a high ortholog prediction (DIOPT) score of 16/19 (Hu et al., 2011). Recent single-cell sequencing data from the adult fly brain indicate that *CG11007* is widely expressed in the adult nervous system in both neurons and glia (Davie et al., 2018). To determine the *in vivo* consequences of loss of *CG11007*, we used the well-established Gal4/UAS system to selectively reduce *CG11007* mRNA in the nervous system in either all neurons or all glia via RNAi (interference). Pan-neuronal knock-down of *CG11007* using nSyb (neuronal Synaptobrevin)-Gal4 at 25°C led to viable flies that were grossly normal. Neuronal reduction of *CG11007* did not cause any significant climbing deficits or mechanically-induced seizures in aged (20-day old flies) flies of either sex at this temperature (Supplemental Figure 7E). Conversely, pan-glial knock-down of *CG11007* using repo-Gal4 led to increased time to climb in aged flies of either sex at 25°C (Figure 7E). Climbing was not significantly different from luciferase RNAi controls in young (5-day old flies). Similarly, glial reduction of *CG11007* at 25°C causes seizure-like behaviour (freezing and/or spasticity) in aged flies after 15 seconds of vortex (Figure 7F). This bang-sensitivity phenotype was observed in both sexes and while not significant, showed a trend in young flies. Our results indicate that the glial function of CG11007 is important for appropriate neuronal function in the organism.

The Gal4/UAS system is temperature-dependent (Nagarkar-Jaiswal et al., 2015) where increased temperature leads to increased Gal4-mediated UAS binding and subsequent RNAi knock-down of the mRNA target. Similar to 25°C, neuronal knock-down of *CG11007* at 29°C led to viable flies that showed no climbing deficits or mechanically-induced seizures in aged (20 days old) flies of either sex at this temperature (Supplemental Figure 7G, H). Interestingly, neuronal knock-down of *CG11007* caused a mild but significant increase in climbing speed in aged flies at 29°C (Supplemental Figure 7H). Regardless, these data indicate that CG11007 is not critical to neuronal function in flies. In contrast, glial knock-down of *CG11007* at 29°C causes sub-viability as flies no longer eclose from pupae at the expected Mendelian ratios (Supplemental Figure 7I). Adult flies that do survive at 29°C have significant climbing deficits (Supplemental Figure 7J) and seizure-like behaviour (Supplemental Figure 7K) at 5 days old when compared to luciferase RNAi controls of the same sex. Lastly, glial knock-down of *CG11007* at 29°C significantly shortens fly lifespan (Figure 7G). 50% survival dropped from 28 days to 18 days in males and from 44 days to 40 days in females. Together, these data indicate the importance of glia TMX2 and CG11007 in long-term health and nervous system function.

## Discussion

Our previous research had identified ER-lumenal ROS production as critical for the increased formation of ERMCS during ER stress (Bassot et al., 2023). However, the formation of ERMCS redox nanodomains on the cytosolic face of these membrane contacts indicates that this location also undergoes ROS-dependent changes (Booth et al., 2016). With the present study, we have identified the ubiquitous TMX2 protein as key in the control of such changes. We have determined that TMX2 cooperates with PRDX1 and to a lesser extent with PRDX2 to control ERMCS cytosolic redox. This result could be tied to the common cytosolic role of thioredoxins as reductases of PRDXs. Alternatively, TMX2 could simply absorb ROS at ERMCS with the help of PRDXs. Regardless, ER membrane-embedded TMX2 therefore connects the ER environment with the cytosol and its redox enzymes. Subsequently, it directs ER and mitochondrial ROS supply away from mitochondrial surface proteins such as TOM70. In the case of this substrate, TMX2 prevents sulfenylation that normally promotes ERMCS tethering. To do so, TMX2 uses a SNDC motif within its thioredoxin domain. This sequence is rare within mammalian thioredoxins but resembles the one found within the cytosolic E. coli chaperedoxin cnoX (SQHC), which cooperates with heat shock proteins to similarly protect its substrates from permanent oxidation (Dupuy et al., 2023, Goemans et al., 2018).

We have therefore identified mitochondrial sulfenylation as a novel signal determining ERMCS spacing: TOM70 sulfenylation correlates with tight ERMCS, while TMX2 prevents this state and promotes wide ERMCS spacing. PRDX1 interaction with TOM70 increases in the former scenario, likely in a futile interaction, but its TOM70 interaction is not detectable in the latter. This indicates that PRDX1 normally eliminates ROS that can otherwise sulfenylate TOM70 in the absence of TMX2. While we used TOM70 as a model substrate in this study, multiple mitochondrial surface proteins are under redox control. Some of these, including VDAC1, are amongst the TMX2 interactome, although in a C170-independent manner. Whether their oxidation state (Reina et al., 2020) plays a role for VDAC-controlled ERMCS functions (Prasad et al., 2015) is unclear.

The consequences of TMX2-mediated ERMCS spacing are wide-ranging for cell metabolism. We detected changes in the mitochondrial generation of cellular energy, and in cellular lipids. Within mitochondria, the OXPHOS electron transport chain undergoes changes in the balance between the individual complexes. Most of these are depleted, while complex II resulted as increased upon interference with TMX2. Interestingly, complex II accepts electrons from the Krebs cycle upon the conversion of succinate to fumarate mediated by SUCLG1/SUCLG2 (Bezawork-Geleta et al., 2017). Given that we observed a significant decrease of SUCLG1 in parallel, our findings indicate a dysregulation of the succinate metabolism with low levels of TMX2. Succinate is thought to act as an oncometabolite, which is associated with altered gene expression, and promotes growth (Selak et al., 2005). Its expected deficiency upon TMX2 interference, resulting in decreased SUCLG1 and increased complex II could therefore have the opposite effect and decrease growth and cell survival. This could explain the association of increased TMX2 with breast cancer progression (Hatzidaki et al., 2020). Similar to this hypothesis, the decreased presence of ether phospholipids, as observed with low levels of TMX2 (Figure 7G), is expected to decrease lifespan (Cedillo et al., 2023) and associated with nervous system defects in patients, including microcephaly and epilepsy (Dorninger et al., 2017). Ether lipids are synthesized in peroxisomes and the ER (Dean & Lodhi, 2018), suggesting TMX2 could impact either location. Likewise, the changes observed with HexCer, a class of ceramide species modified with galactose or glucose, are another cause of mitochondrial ROS production due to the impaired activity of ceramide transfer protein at the ER (Rao et al., 2014).

From the characterization of the redox and metabolic phenotype, we would predict that rescuing succinate and ether phospholipid levels could provide a therapeutic avenue for the treatment of NEDMCMS. To explore such possibilities, we could use our *in vivo* model, where we detected a reproduction of the clinical features of this disease, comprising defective brain function and increased epileptic readouts. Importantly, these defects were only manifest if TMX2 was reduced within glial cells. Due to the similarity of NEDMCMS with other developmental epileptic encephalopathies, our findings may extend to those diseases as well. For example, the ERMCS regulatory protein PACS-2 is mutated in developmental and epileptic encephalopathy-66 (DEE66 – MIM#618067) (Olson et al., 2018). Recent evidence showed that DEE66 results from increased availability of Ca^2+^ in the cytosol, which then activates synaptic vesicle release in neurons (Thi My Nhung et al., 2023). It remains to be shown whether the two diseases share a common mechanistic basis. Alternatively, the increased mitochondrial activity upon TMX2 interference could compromise normal levels of astrocytic lactate production, which is necessary for neuronal survival (Demetrius et al., 2014). Within this cell type, TMX2 could be part of the antioxidant response. This response is dependent on Nrf2, which is less active in neurons (Kraft et al., 2004). Alternatively, nuclear factor of activated T cells 1 (NFAT1) is known to induce ER antioxidant proteins including TMX1, TMX2 and GPx8 (Mognol et al., 2019) and promotes glial activation (Furman & Norris, 2014). Further experiments will have to determine whether Nrf2 or NFAT-1 control TMX2. Such insight could then be used to explore therapeutic approaches with the aim to rescue TMX2 and/or PRDX function. Any such approach should then restore neuronal Ca^2+^ availability that controls cell death and synaptic activity, as well as the metabolic pathology in glial cells.

### Limitations of the Study

While our manuscript delineates the function of TMX2 for the formation of ERMCS, it currently does not clearly describe the biochemical activity of TMX2 for PRDX1. We provide data that demonstrate a partial replication of TMX2 depletion in NEDMCMS patient cells, but which function is critical for the pathology is unclear. The identification of glial cells as critical in our fly model needs to be validated in NEDMCMS.

## Author contributions

Conceptualization: JC & TS; Data curation: JC, MCY, AB, DMP, TM, JZ, HM, & SGF; Formal analysis: JC, MCY, DMP, TM, JZ, & SGF; Funding acquisition: PCM & TS; Investigation: JC, MCY, AB, DMP, TM, JZ, HM, RB, SGF, YF, AZB & JM; Methodology: JC, MCY, AB, DMP, TM, JZ, PCM & TS; Project administration: TS; Resources: KB, LL, MO, MJL, HL, WHT, GMSM, BM, PCM & TS; Software: JC, AB, & TM; Supervision: PCM & TS; Validation: PCM & TS; Visualization: JC & MCY; Roles/Writing - original draft: TS; Writing - review & editing: JC, MJL, BM, PCM & TS.

## Acknowledgments

We thank Tjakko J. van Ham, Peter Hwang and Leslie Poole for many helpful suggestions. We thank the following colleagues for critical reagents: Tito Cali for the SPLICS constructs, and Jennifer Rieusset for FATE-1. We are indebted to Xuejun Sun, Priscilla Gao and the Cell Imaging Facility at the Cross Cancer Institute/University of Alberta for assistance with microscopy. Funding for this study has been provided by CIHR operating grant PS162449 and CRS grant 834492 to TS. PCM is supported by operational funds from NSERC (Discovery Grant RGPIN-2023-05422). DMP is supported by the CIHR Canada Graduate Scholarship (CGS-M).

## Declaration of Interests

The authors declare no competing interests.

## METHODS

### EXPERIMENTAL MODEL AND STUDY PARTICIPANT DETAILS

U251 MG cells (ATCC, US), HeLa cells (ECACC, UK), and HEK293 cells (ECACC, UK) were cultured in DMEM (Gibco) with 10% fetal bovine serum (FBS; F1051, Sigma-Aldrich) at 37°C under 5% CO_2_. Patient-derived skin fibroblasts were provided by Dr. Grazia M.S. Mancini’s lab (Erasmus University Medical Center, Netherlands) and Dr. Wen-Hann Tan’s lab (Harvard Medical School, US), which were grown under the conditions previously described (Vandervore et al., 2019).

All flies were raised on standard molasses-based food at 22 °C unless otherwise indicated. The *nSyb-Gal4* and *repo-Gal4* stocks were obtained from the Bloomington Drosophila Stock Center (BDSC) using the following genotypes: *y^1^ w*; P[nSyb-GAL4.S]3* (BDSC_51635) and *w^1118^; P[GAL4]repo/TM3,Sb1* (BDSC_7415). The *UAS-luciferase RNAi* was described previously (Marcogliese et al., 2018). The *UAS-CG11007 RNAi* (*w^1118^; P[GD2093]v40833*) was obtained from the Vienna Drosophila Resource Center.

### METHOD DETAILS

#### DNA constructs and small interfering RNA (siRNA)

The expression plasmids for C-terminally HA-tagged human TOM70, human Flag-TMX2 (N-terminus, pcDNA3.1, wild-type and all mutants); human Flag-TMX2 (at 49 aa, pcDNA5, WT), and human Flag-TMX2 (pIRES2-EGFP) were constructed by Genscript (Piscataway, NJ). Dog Flag-Calnexin was previously described (Lynes et al., 2012). Plasmid-based transfection was performed with Lipofectamine 3000 (Invitrogen) for U251 cells, and Metafectene Pro (Biontex) for HeLa and HEK293 cells, following the manufacturer’s protocols. Cells were processed for experiments 24-48 h after transfection.

For gene silencing, cells were transfected with the Stealth™ RNAi (12935300, Med GC) as a negative control and siRNA (Invitrogen) to silence TMX2 or TOM70 expressions (siTMX2-1, TMX2HSS121596, 5′-CCUAUCUAUGCUGACCUCUCCCUUA-3′; siTMX2-2, TMX2HSS181843, 5′-GGGAAGGUGGAUGUUGGACGCUAUA-3′; siTOM70-1, TOM70HSS114910, 5′-GAAAUAGAUGCUGAAGGCAAAUACA-3′; siTOM70-2, TOM70HSS114911, 5′-UGCUGUUGCAUGUACAUGCUGCCUC-3′; siTOM70-3, TOM70HSS114912, 5′-AUAUGUUGUAGCAUUAUCUGGUUCC-3′;). Following the manufacturer’s recommendations, RNA interference was performed using Oligofectamine (Invitrogen). Cells were collected for experiments 48-72 h after transfection.

#### TMX2 CRISPR/Cas9 knockout

U251 TMX2 KO cells were generated with the TMX2 Human Gene Knockout Kit (OriGene, KN400032). The pCas-Guide vector (OriGene, GE100003) with a scrambled sequence was used as a control. Briefly, U251 MG cells were co-transfected with a pCas-Guide vector (target sequence: CCAACCTTACTACTACCTTCTGT) and a linear donor vector (containing EF1a-GFP-P1A-Puro) using Lipofectamine 3000. Cells were grown for two weeks before the selection of puromycin (P8833, Sigma-Aldrich). Then, the cells were transferred to 15-cm dishes, and individual cell colonies were collected after culturing with 0.25 µg/mL puromycin for three weeks. The insertion of the donor vector was verified by the presence of a GFP signal via flow cytometry, and TMX2 knockout was confirmed by Western blot.

#### Stable cell lines generation

U251 cells were transfected with pIRES2-EGFP plasmids (empty vector, TMX2-WT, TMX2-C170S, and TMX2-S167C) using Lipofectamine 3000. 24 h after transfection, cells were cultured with 1 mg/ml of Geneticin (10131027, Thermo Fisher Scientific). After 2 weeks, individual colonies were selected and verified with Western blot and flow cytometry.

#### Cell lysates and immunoblotting

After washing 2 times with ice-cold 1× PBS (14080-055, Gibco), cells were trypsinized or scrapped with a rubber policeman. Cells were then resuspended in m-RIPA lysis buffer [50 mM Tris-HCl (pH 7.4), 150 mM NaCl, 5 mM MgCl_2_, 1% NP-40, 0.25% Sodium deoxycholate] containing 1× cOmplete EDTA free protease inhibitor (11873580001, Roche) and 1× PhosSTOP phosphatase inhibitor (4906845001, Roche), incubating on ice for 10 min. The lysates were then vortexed for 5 s and centrifuged at 12,000 g for 15 min at 4°C to get post-nuclear supernatant. The protein concentration of lysates was tested by the Pierce BCA Protein Assay Kit (23225, Thermo Fisher Scientific). For the reducing gel, cell lysates were mixed with the sample loading buffer [60 mM Tris-HCl (pH 6.8), 2% SDS, 10% glycerol, 5% β-mercaptoethanol, and 0.004% bromophenol blue] and boiled at 75°C for 8 min. For the non-reducing gel, the lysates were supplemented with 20 mM *N*-ethylmaleimide (NEM; E3876, Sigma-Aldrich) and mixed with loading buffer without β-mercaptoethanol. The equal amounts of protein per sample were loaded onto sodium dodecyl sulfate-polyacrylamide gel electrophoresis (SDS-PAGE) gels. The gels were electrophoresed in the running buffer [25 mM Tris-HCl (pH 7.3), 200 mM glycine, 0.1% SDS] for 75 min at 150 V, followed by transfer to nitrocellulose membranes (Bio-Rad) for 2 h at 400 mA. Then the membranes were blocked with 2% skimmed milk in TBST [10 mM Tris-HCL (pH 8.0), 150 mM NaCl, 0.05% Triton X-100 (1086431000, Sigma-Aldrich)] for at least 30 min and then kept with the primary antibodies overnight at 4°C. After washing 3 times with TBST, the membranes were probed with secondary antibodies for 1 h. Then the membranes were washed with TBST and scanned using an LI-COR Imager (CLX-2901, Biosciences). Image Lab Software (Bio-Rad) intensity was used for band intensity quantification and analysis.

#### Protease K protection assay

U251 MG cells were transfected with two versions of Flag-TMX2-wild-type plasmids. The Flag-tag was inserted following the putative signal peptide (Version 1) or at the N-terminus (Version 2). Cells were harvested with homogenization buffer [20 mM HEPES (pH 7.4), 250 mM sucrose, 1 mM EDTA, 1 mM EGTA] in the presence of 1× protease inhibitor, followed by homogenization with a ball-bearing homogenizer (ball clearance 14 µm, Isobiotec). Next, the lysates were spun at 2,000 rpm for 30 min at 4°C, using a JA-12 rotor (Beckman) to remove nuclei and unbroken cells. The post-nuclear supernatants were centrifuged at 60,000 rpm for 60 min at 4°C with a TLA 120.2 rotor (Beckman) to collect ER-mitochondria-enriched fractions. Pellets were resuspended in homogenization buffer, divided equally into tubes, and treated with proteinase K (P8107, NE BioLabs) at different concentrations for 10 min at room temperature (R.T.). The reaction was quenched by incubating with 4 mM phenylmethylsulfonyl fluoride (PMSF; 52332, Sigma-Aldrich) for 5 min on ice. Then the samples were boiled at 75°C for 10 min in the loading buffer. All samples were sonicated and processed for SDS-PAGE and immunoblotting.

#### Percoll gradient fractionation

HeLa and U251 MG cells were seeded in 15 cm dishes, collected with 5 ml of homogenization buffer containing 1× protease inhibitor, and spun at 1,500 rpm for 5 min in a JA-12 rotor. Then cells were resuspended in 5 ml homogenization buffer and passed 15-25 times through the homogenizer (ball clearance 14 µm). The homogenates were centrifuged thrice at 2,000 rpm to remove unbroken cells and nuclei. The supernatant was saved as a fraction of homogenate. The remaining supernatants were centrifuged at 8,500 rpm for 10 min to obtain crude mitochondria fractions. Next, the resulting supernatants were spun at 60,000 rpm for 60 min at 4°C in a TLA 120.2 rotor to acquire the fractions of microsome (pellet) and cytosol (supernatant, incubation with ice-cold 100% acetone overnight). The previously isolated crude mitochondria fractions were layered on the 18% Percoll gradient (GE17-0891-02, Sigma-Aldrich) in OptiSeal polypropylene tubes (361623, Beckman). Then the samples were spun at 33,300 rpm for 30 min with a 90Ti rotor (Beckman) to get the MAM and pure mitochondria fractions. To remove the Percoll, the MAM fractions were spun at 60,000 rpm for 1 h using the TLA 120.2 rotor. The fractions of pure mitochondria were transferred to tubes supplemented with homogenization buffer and centrifugated twice at 10,000 g for 10 min, and the pellets were saved. Equal amounts of samples were analyzed by immunoblotting, as previously described (Wieckowski et al., 2009).

#### Mitochondria isolation

Mitochondria were isolated from wild-type and TMX2 KO cells using the Mitochondria Isolation Kit for Cultured Cells (89874, Thermo Fisher Scientific) according to the manufacturer’s protocol (Option A). Briefly, cells were harvested with PBS and centrifuged at 850 g for 2 min to obtain cell pellets. 800 µL of Reagent A was added to each pellet, vortexed for 5 s at medium speed, and incubated on ice for 2 min. Then the homogenate was added 10 µL of Reagent B and vortexed every minute at maximum speed. Next, 800 µL of Reagent C was added to each tube and inverted multiple times to mix. After centrifuging at 700 g for 10 min, the mitochondria-enriched supernatants were transferred to new tubes and spun again at 12,000 g for 15 min at 4°C. Lastly, the isolated mitochondria pellets were washed with 500 µL Reagent C, resuspended in homogenization buffer, and maintained on ice before downstream experiments.

#### Glycosidase Digestion

The experiments were performed with U251 MG cells to test endogenous TMX2 and HeLa cells to test TMX2 putative glycosylation mutants. On the experiment day, cells were lysed with glycoprotein denaturing buffer (B1704S, NE BioLabs). Each sample was sonicated, denatured at 100°C for 10 min, cooled on ice, and centrifuged for 10 s. The lysates were then treated with 500 U of peptide-N-glycosidase F (PNGase F; P0704L, NE BioLabs) in a reaction solution containing 50 mM sodium phosphate (pH 7.5) and 1% NP-40 (B2704S, NE BioLabs) for 1 h at 37°C. For endoglycosidase H (Endo H; P0702S, NE BioLabs) digestion experiments, the cell lysates were deglycosylated with 500 U of Endo

H in the reaction buffer containing GlycoBuffer 3 (B1720S, NE BioLabs) for 1 h at 37°C. Then equal amounts of samples were analyzed by immunoblotting, as described previously (Qiu et al., 2023).

#### Immunoprecipitation

For cross-linking immunoprecipitation, cells were incubated with 2 mM dithiobis(succinimidyl propionate) (DSP; 22585, Thermo Fisher Scientific) in 1× PBS++(containing CaCl_2_ and MgCl_2_, diluted from 10× PBS++, 1014200-075, Gibco) for 30 min at R.T. Next, cells were washed twice with PBS++ and quenched with 10 mM of NH_4_Cl (A9434, Sigma-Aldrich) in PBS++ for 10 min at R.T. The cells were then washed with PBS++ and harvested in RIPA lysis buffer. For co-immunoprecipitation, cells were washed with PBS++ and collected in CHAPS lysis buffer [10 mM Tris (pH 7.4), 150 mM NaCl, 1 mM EDTA, 1% CHAPS] containing 20 mM NEM. Then the lysates were rocked for 30 min at 4°C and centrifuged at 12,000 g for 15 min to pellet nuclei and unbroken cells. Preclearing was performed by incubation with Protein G Magnetic Beads (10004D, Invitrogen) for 1 h at 4°C. Then the indicated antibodies were added to the supernatants and rocked overnight at 4°C. Protein G Magnetic Beads were added and rocked for 1 h at 4°C. The beads were washed 4 times with RIPA or CHAPS buffer, boiled in the sample loading buffer at 75°C for 10 min, and analyzed by immunoblotting (Bassot et al., 2023).

#### DCP-Bio1 assay for identification of sulfenylated proteins

DCP-Bio1, consisting of the sulfenic acid-reactive 3-(2,4-dioxocyclohexyl) propyl (DCP) linked to biotin, was used to capture the sulfenic acid formation (NS1226, Sigma-Aldrich) (Klomsiri et al., 2010, Nelson et al., 2010). Briefly, HEK293 cells were cultured in 10-cm dishes and transfected with the indicated RNAi for 48 h. After washing twice with PBS++ (containing 10 mM NEM), cells were lysed in the reaction buffer [50 mM Tris-HCl (pH 7.5), 100 mM Sodium Chloride, 0.1% SDS, 0.5% Sodium Deoxycholate, 0.5% NP-40, 0.5% Triton X-100] supplemented with 1 mM DCP-Bio1, 100 µM Diethylene triamine pentaacetic acid (DTPA), 10 mM NEM, 1 mM PMSF, 200 U/mL catalase, and 1× protease inhibitor. Samples were kept on ice for 30 min, followed by centrifugation at 16,000 g for 10 min at 4°C. Then the post-nuclear supernatants were kept on ice for 90 min in the dark for labeling of the sulfenic acids. Zeba™ Spin Desalting columns (89890, Thermo Fisher Scientific) were pre-washed 3 times with CHAPS lysis buffer, loaded with the reacted samples, and spun at 1,000 g for 4 min at 4°C to remove unreacted DCP-Bio1. Then protease inhibitor was added back to the lysates, and protein concentration was determined by the BCA assay as described. Lysates were incubated with streptavidin beads (65305, Invitrogen) overnight at 4°C. The beads were washed three times with CHAPS lysis buffer and boiled in the sample loading buffer supplemented with 50 mM dithiothreitol (DTT; A22066A, Invitrogen) for 10 min at 75°C. The lysates were analyzed by SDS-PAGE and Western blot.

#### *In vitro* roGFP2 measurements

YPH499 Δ*glr1*Δ*grx1*Δ*grx2* cells were co-transformed with a p415TEF plasmid encoding the *Saccharomyces cerevisiae* plasma membrane glutathione transporter Opt1/Hgt1 and a p416TEF plasmid encoding a genetic fusion construct consisting of roGFP2 and HsGrx1, *Hs*TMX2 or mutants thereof with a short interspacing polypeptide linker (SGGGG × 5). Cells were grown to late logarithmic phase, *OD*_600_

≈ 3.0, in Hartwell’s complete medium with 2% glucose and lacking leucine and uracil to facilitate plasmid retention. For fluorescence measurements, cells were harvested at 1000 g for 3 min at R.T. and resuspended in 100 mM MES/Tris buffer (pH=6) to an *OD*_600_ = 7.5. Subsequently, 200 µl aliquots were transferred to the well of a flat-bottomed 96-well microtitreplate. The plate was centrifuged at 30 g for 5 min at R.T. so that the cells formed a loose pellet at the bottom of each well. The measurement was initiated by the addition of H_2_O_2_ or GSSG at the indicated concentration and roGFP2 oxidation was followed for another 30 min in a BMG Labtech CLARIOstar^Plus^ fluorescence plate reader.

RoGFP2 contains two redox-active cysteine residues on opposing β-strands adjacent to the chromophore (Zimmermann et al., 2021). The formation of a disulfide bond between these cysteine residues results in a ratiometric change in the fluorescence excitation spectrum of the roGFP2. The neutral chromophore, which is predominant in disulfide bond-containing roGFP2, shows an excitation maximum at ∼400 nm, while the anionic chromophore, which is predominant in reduced roGFP2, is excited with a maximum of ∼490 nm. Excitation at both 400 nm and 490 nm results in fluorescence emission with a maximum of ∼510 nm, therefore, allowing ratiometric measurements. For roGFP2 calibration, 20 mM N, N-dimethylformamide (Diamide; D3648, Sigma-Aldrich), or 100 mM DTT were added to the corresponding wells to obtain fully oxidized or fully reduced control samples. Probe calibration allows for the determination of the degree of roGFP2 oxidation (OxD) according to the equation below:

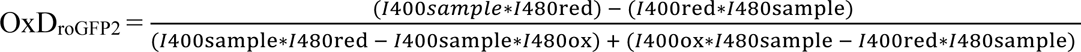

#### Mitochondrial Ca^2+^ measurements

Live-cell imaging was performed with an Olympus FV1000 laser-scanning confocal microscope using a 20× objective (XLUMPLANFL, NA 1.0, Olympus) (Gutierrez et al., 2020). The microscope was equipped with a megapixel camera (PL-A686 6.6, PixeLink) and a perfusion system with a peristaltic pump (Watson-Marlow Alitea-AB, Sin-Can). The flow speed was set at 6 ml per minute. Images were obtained with the Olympus FluoView software and then analyzed with Fiji (NIH, USA) using the Time Series Analyzer plugin (version 3.0).

The low-affinity Mito R-GECO fluorescent probe was used to measure mitochondrial Ca^2+^. To measure ER-mitochondria Ca^2+^ transfer, cells were cultured on the poly-L-lysine (P4707, Sigma-Aldrich) coated 12-mm coverslips (Fisher Scientific) and transfected with the fluorescent plasmid. 24 h after transfection, coverslips were transferred to the microscope chamber, perfusing with Hanks’ balanced salt solution (HBSS) with Ca^2+^ and Mg^2+^ (HBSS/Ca^2+^/Mg^2+^; 14025092, Thermo Fisher Scientific). After establishing a baseline for 30 s, the perfusion buffer was replaced by HBSS/Ca^2+^/Mg^2+^ supplemented with 50 mM histamine (H7250, Sigma-Aldrich) for 5 min. To measure mitochondrial Ca^2+^ content, the buffer was supplemented with 10 µM FCCP (15218, Cayman) for 15 min after establishing a fluorescence baseline. Images were captured every second under histamine treatment and every 3 sec under FCCP treatment in the Alexa 546 channel (559 nm excitation/575-675 nm emission).

#### ER Ca^2+^ measurements

ER R-GECO fluorescent probe was used to quantify ER Ca^2+^ content. Cells were prepared as mentioned above. After the first 30 s, the buffer was replaced with HBSS/Ca^2+^ free/Mg^2+^ free (14175095, Thermo Fisher Scientific) containing 60 µM tert-butylhydroquinone (TBHQ; 112941, Sigma-Aldrich), 410 µM MgSO_4_ (230391, Sigma-Aldrich), 1.75 µM MgCl_2_ (MX0045-1, EM Science), and 100 µM EGTA (4100, OmniPur EMD) for 15 min to inhibit the activity of SERCA pump. Live-cell images were captured every 3 sec. As described previously, to measure Ca^2+^ release from ER, cells were perfused with 50 mM histamine for 5 min, and images were captured every second in the Alexa 546 channel (559 nm excitation/575-675 nm emission).

#### High-resolution respirometry

Cells were seeded in 6-well at least 48 h before experiments. The Oxygraph-2k machines (Oroboros Instruments, Innsbruck, Austria) were washed with 70% ethanol, 90% ethanol, and Milli-Q water. Then the oxygen was calibrated at 37°C, and the chambers were filled with 2 mL ice-cold DMEM, according to the manufacturer’s instructions. For respirometry measurements in intact cells, cells were trypsinized and neutralized in fresh DMEM, followed by centrifugation at 350 g for 5 min at R.T. The cells were then resuspended in 2.6 mL of DMEM, and cell number was calculated. 2 mL of the intact cells were loaded into each chamber, and cellular respiration was recorded by measuring the oxygen flux. Briefly, basal respiration was measured without any added substrates. 2 µg/mL oligomycin (141829, Abcam) was added to inhibit ATP synthase and estimate the leak respiration. Maximal respiration was stimulated by the stepwise addition of the uncoupler carbonyl cyanide 4-(trifluoromethoxy)phenylhydrazone (FCCP; C2920, Sigma-Aldrich). Lastly, 0.25 µM antimycin A (A8674, Sigma-Aldrich) was added to inhibit Complex III and measure non-mitochondrial respiration.

For permeabilized cell respiration, cells were counted and diluted in 2.6 mL of MiR05 buffer [110 mM D-sucrose, 60 mM potassium lactobionate, 20 mM taurine, 20 mM HEPES, 10 mM KH_2_PO_4_, 3 mM MgCl_2_·6H_2_O, 0.5 mM EGTA and 1 g/L fatty acid-free BSA (10775835001, Sigma-Aldrich), at pH 7.1]. The digitonin concentration used for permeabilization was optimized before experiments. The following pathways were measured: NADH pathway (N-pathway, NADH-linked substrates, such as pyruvate and malate entering the chain at complex I) was measured after the addition of 5 mM pyruvate (P2256, Sigma-Aldrich) + 2 mM malate (M0875, Sigma-Aldrich) + 10 mM glutamate (G1626, Sigma-Aldrich) and 2.5 mM ADP (A5285, Sigma-Aldrich). NADH-Succinate pathway (NS-pathway, convergent electron flow at the Q-junction through Complexes I+II) was measured by the addition of 10 mM succinate (S2378, Sigma-Aldrich) (Mast et al., 2022). The succinate pathway (S-pathway; succinate entering the chain through complex II) was tested by inhibiting complex I with 0.5 µM rotenone (R8875, Sigma-Aldrich). After adding 0.5 mM tetramethyl phenylenediamine (TMPD; T3134, Sigma-Aldrich) and 2 mM ascorbate (A4034, Sigma-Aldrich), Complex IV activity was measured. The reduced state of TMPD was maintained by ascorbate to feed electrons into complex IV. Then the chemical background was subtracted after adding 100 mM sodium azide (S2002, Sigma-Aldrich). Mitochondrial respiration was determined by O_2_ flux per cell and Flux Control Ratio (normalized to the maximal respiration). Datlab7 software (Oroboros Instruments) was used for data acquirement and analysis.

#### Immunofluorescence microscopy

For standard immunofluorescence, cells were cultured on the poly-L-lysine-coated coverslips (18 mm, thickness #1.5) in 6-well plates, as previously described. On the experiment day (24 or 48 h after transfection), cells were rinsed 3 times with PBS++ and fixed with 2% paraformaldehyde (PFA; 15710, Electron Microscopy Sciences) in PBS++ for 15 min at R.T. Then the cells were rinsed 3 times with PBS++ to quickly remove most of the fixative and quenched with 50 mM NH_4_Cl for 10 min. Next, cells were washed with PBS++, permeabilized with 0.1% Triton X-100 in PBS++ for 10 min, and rinsed with PBS++ before blocking with 2% BSA in PBS++ for 1 h. Cells were then incubated with primary antibodies [rabbit anti-TMX2 (1:100, 19838-1-AP, Proteintech); mouse anti-TOM70 (1:150, H00009868-B01P, Abnova); mouse anti-Erp57 (1:100, SMC-168B, StressMarq); mouse anti-Flag (1:200, 200-301-B13, Rockland); rabbit anti-Flag (1:400, 14793, Cell Signaling); mouse anti-HA (1:200, 901513, BioLegend)] for 1-2 h at R.T. in a humidified, light-protected chamber. After washing 3 times with PBS++ with 5 min rocking, cells were incubated with secondary antibodies [goat anti-Rabbit IgG (H+L), Alexa Fluor™ 594, A-11012, Invitrogen; goat anti-Mouse IgG2a, Alexa Fluor™ 488, A-11029, Invitrogen, at 1:1000], for at least 30 min. Cells were washed 3 times with PBS++, and coverslips were then mounted cell side down on the slides with ProLong™ Gold Antifade Mountant (P36934, Invitrogen). For mitochondrial morphology, cells were treated with 100 nM MitoTracker^TM^ Red CMXRos probe (M7512, Thermo Fisher Scientific) for 30 min in the incubator before fixation. Nuclei were stained with 4′,6-diamidino-2-phenylindole dihydrochloride (DAPI; D9542, Sigma-Aldrich) for 5 min at 1:1000.

The split-GFP-based contact site sensor (SPLICS) was designed to emit fluorescence when the ER and mitochondria are in proximity (Cali & Brini, 2021, Cieri et al., 2017). For co-localization analyses, two versions of SPLICS were used: SPLICS-short measures the organelle distances between 8 and 10 nm, and SPLICS-long measures their contacts between 40 and 50 nm. HeLa cells were transfected with SPLICS-long or SPLICS-short plasmids. The next day, cells were cultured on the coverslips (18 mm) and immunostained, as mentioned previously, including fixation in 2% PFA for 15 min and blocking in 2% BSA in PBS++. Mander’s coefficient was calculated on raw images after background subtraction using ImageJ software (NIH, USA).

*In situ* proximity ligation assay (PLA) was performed to detect, visualize, and quantify protein interactions (within 40 nm) as a single dot (Tubbs et al., 2014). 24 h after gene silencing or plasmid DNA transfection, HeLa cells were cultured on the coverslips (18-mm). The next day, cells were fixed and incubated with anti-IP_3_R1 (1:200, 3809289, Millipore) and VDAC1 (1:200, 14734, Abcam) or HA primary antibodies in a humidified chamber at 4°C overnight. Then the cells were washed twice with TBST, followed by incubating with the secondary antibodies (anti-Mouse MINUS, DU092004, Sigma-Aldrich; anti-Rabbit PLUS, DUO92002, Sigma-Aldrich) for 1 h at 37°C. The proximity ligation and polymerase amplification steps followed the manufacturer’s protocols (Duolink^®^ In Situ Detection Reagents Red, DU092008, Sigma-Aldrich). Note that the 5× ligase and 5× polymerase were diluted and added immediately before addition to the samples. Then the cells were stained with DAPI and actin (1:1000, 176753, Abcam). Images were deconvolved, and the number of dots per nucleus was quantified with ImageJ software as described previously (Tubbs & Rieusset, 2016). Experiments were repeated twice, and at least 8 fields were taken in each condition.

Images were obtained with a Leica SP8 Falcon microscope (100× 1.4 NA oil immersion lens). The Leica TCS SP8 microscope was equipped with the Leica Application Suite X Software (version 3.5). Between 4 and 8 z-stack images were captured with 200-400 nm step size. 488 nm excitation was used for SPLICS fluorescence and Alexa 488-labeled antibodies. 594 nm excitation was used for MitoTracker^TM^, PLA dots, and Alexa 594-labeled antibodies. A 405 nm laser was used for DAPI excitation. After background signal subtraction, maximum intensity projection was selected to acquire the summed intensities.

#### Transmission electronic microscopy (TEM)

After gene silencing or plasmid DNA transfection, cells were seeded on the ACLAR^®^ Film (10501, TED PELLA, INC) to 80% confluence. Cell monolayers were fixed with 2% PFA and 2.5% glutaraldehyde (GTA; G5882, Sigma-Aldrich) in 0.15 M cacodylate solution (C0250, Sigma-Aldrich) with 2 mM CaCl_2_ for 20 min at 37°C and 40 min at R.T. Post fixation was done with 2% osmium tetroxide (OsO_4_; 201030, Sigma-Aldrich) in 1.5% potassium ferrocyanide with 2 mM CaCl_2_ (freshly prepared before use) for 1 h on ice. Then the samples were treated with filtered thiocarbohydrazide (TCH; 223220, Sigma-Aldrich) solution for 20 min at R.T., 2% OsO_4_ for 30 min at R.T., and washed with ddH_2_O before staining with 1% uranyl acetate overnight at 4°C on a rocker. The next day, the samples were stained with Lead Nitrate in the aspartic acid solution for 30 min at 60°C. Then the samples were dehydrated with increasing concentrations of ice-cold ethanol (20%, 50%, 70%, and 90% ethanol 5 min each, 100% ethanol 5 min twice) and infiltrated with acetone for 10 min × 2 at R.T. Samples were embedded with Durcupan™ ACM (44611, Sigma-Aldrich) and polymerized at 60°C oven for 48 h before sectioning. A Transmission Electron Microscope (TEM) from JEOL (JEM-2100) equipped with a Lab6 filament at 200 kV acceleration voltage was used for TEM observation of the samples. The system is operated with a Gatan GIF Quantum energy filter using a 2k × 2k digital camera with DigitalMicrograph software (version 2.0). The images were taken with a resolution of 2048 × 2048 pixels. Images were quantitatively analyzed for ER and mitochondria contacts (within 50 nm) with imageJ, as described previously (Bassot et al., 2023, Theurey et al., 2016). Data was generated from at least 30 images per condition with around 100 measurements each.

#### Cellular ATP measurements

Cells were cultured in 12-well plates and reached 80% confluence on the experiment day. Cells were incubated with or without 3 µM oligomycin to estimate mitochondrial ATP production for 1.5 h. Cells were then trypsinized and resuspended in the CHAPS lysis buffer containing 1× protease inhibitor. Cellular ATP content was measured by the ATP Determination kit (A22066, Molecular Probes) following the manufacturer’s instructions, which measured ATP by recombinant the firefly luciferase and its substrate D-luciferin. Then the luminescence (emission maximum 560 nm at pH 7.8) was monitored with a Synergy 4 plate reader (Bio Tek), and the ATP levels were normalized to protein concentrations (Gutierrez et al., 2020).

#### Flow cytometry

Cytosolic ROS, mitochondrial ROS, and mitochondrial membrane potential were measured using a BD LSR Fortessa-SORP flow cytometer (BD Biosciences). Briefly, cells were incubated with 0.5 µM CellROX™ Orange (C10493A, Invitrogen) for 1 h or 1 µM MitoSOX™ (M36008, Invitrogen) for 40 min at 37°C. Then the cells were collected and resuspended in 300 µL HBSS/Ca^2+^ free/Mg^2+^ free supplemented with 0.5% bovine serum albumin (BSA; A6003, Sigma-Aldrich). Single, live cells of interest were gated on the forward scatter (FSC) and side scatter (SSC) plots. The gates were determined by control groups, treating cells with either 400 µM tert-butyl hydroperoxide (TBHP; C10493A, Invitrogen) or 5 mM N-acetyl-L-cysteine (NAC; A9165, Sigma-Aldrich) for 1 h at 37°C. Then the fluorescence intensity of CellROX™ Orange (545 nm excitation/565 nm emission) and MitoSOX™ (510 nm excitation/ 580 nm emission) were measured. Mitochondrial membrane potential was tested after incubating cells with 40 nM tetramethylrhodamine, methyl ester (TMRM, T668, Invitrogen) for 30 min at 37°C. The gates were adjusted by treating cells with 3 µM oligomycin or 10 µM FCCP for 1.5 h. Then the TMRM fluorescence intensity was measured (552 nm excitation/579 nm emission). Mean fluorescence intensity was determined, and data analysis was performed with the BD FACSDiva™ software.

#### The RNA extraction and RT-qPCR

The total RNA was extracted using TRIzol (Life Technologies) following the manufacturer’s protocol. Two micrograms of total RNA were subjected to reverse transcription using the Superscript II kit (Life Technologies) and the random primers (Life Technologies). The relative gene abundance in the cDNA sample was examined by the quantitative PCR (qPCR) experiment on Mastercycler Realplex^2^ (Eppendorf) using PerfeCTa SYBR green PCR mix (Quanta Bioscience). Oligonucleotides for the reaction were summarized in Table X. The change in the mRNA level of the genes among various mutants was evaluated by the comparative C_T_ method (Schmittgen & Livak, 2008). The C_T_ value for each gene target was normalized against the internal control (ACTB) to give the ΔC_T_ value. The ΔΔC_T_ for each gene was calculated as a difference in ΔC_T_ values between the wild-type and the TMX2-KO cells. The difference in gene mRNA levels between wild-type and TMX2-KO cells was given as 2^-ΔΔCT^, based on the assumption that the amplification efficiency of the PCR reaction is 100%.

#### Liquid chromatography-mass spectrometry (LC-MS)

MAM and mitochondria proteomics altered in the absence of TMX2 were quantified by LC-MS. MAM and mitochondria samples were isolated as described above. The isolated mitochondria pellets were incubated with or without 40 µg/ml proteinase K for 30 min at R.T. The reaction was stopped by adding 4 mM PMSF for 5 min on ice. Samples were sonicated, and 50 µg of each was loaded and ran on the precast gels (4561034, Bio-Rad), stained with Coomassie blue P-250 (161-0436, Bio-Rad), and started the in-gel digestion process (Rashed et al., 2021). Briefly, gel fragments were destained twice in the ammonium bicarbonate/acetonitrile (50:50) solution. Then the samples were reduced with DTT and alkylated with iodoacetamide, followed by digestion with the sequencing grade modified trypsin (V5111, Promega) overnight at 37°C. Tryptic peptides were extracted from the gel with 97% water, 2% acetonitrile (047138-M1, Thermo Fisher Scientific), 1% formic acid (270480025, Thermo Fisher Scientific), and then a further extraction with the previous extraction buffer and acetonitrile (50:50). For the IP-MS, proteins were cross-linked with 2 mM DSP as described previously and immunoprecipitated using Anti-Flag^®^ M2 Magnetic Beads (M8823, Sigma-Aldrich) or protein targeted primary antibodies and Protein G Magnetic Beads. Trypsin digestions were performed on a KingFisher™ Duo Prime Purification System (Thermo Fisher Scientific). Samples were reduced and alkylated (DTT, iodoacetamide) before digestion with the sequencing grade modified trypsin for 10 h at 37°C.

The tryptic peptides were resolved with the EASY-nLC 1000 (Thermo Fisher Scientific) connected to the Q-Exactive Orbitrap mass spectrometer (Thermo Fisher Scientific) with an EASY-Spray™ capillary HPLC column (75 µm × 25 cm, 100Å, 2 µm; ES902A, Thermo Fisher Scientific). The mass spectrometer was operated in data-dependent acquisition (DDA) mode with a m/z range of 300-1700 and 35,000 resolution. By using HCD dissociation, the 12 most intense multiply charged ions were sequentially fragmented, and their fragments spectra were recorded in the orbitrap at 17,500 resolution. For identification of the sulfenylated cysteines, peptides were resolved using an EASY-Spray capillary HPLC column (ES902A, 75 µm × 25 µm, 100Å, 2μm, Thermo Scientific) and a Vanquish Neo UHPLC (Thermo Fisher Scientific) coupled to an Orbitrap Exploris 480 mass spectrometer (Thermo Scientific). The mass spectrometer was operated in data-dependent acquisition mode with a resolution of 60,000, an m/z range of 350–1700, and a cycle time of 3 seconds. Selected ions were fragmented by HCD dissociation, and spectra of their fragments were recorded in the orbitrap at a resolution of 15,000. After fragmentation, all precursors selected for dissociation were dynamically excluded for 30 s.

Data was analyzed with Proteome Discoverer 2.5 (Thermo Fisher Scientific), and a human proteome database (UniProt) was screened with SEQUEST (Thermo Fisher Scientific). The normalization node was added to the processing workflow. A false discovery rate (FDR) between 0.01 and 0.05, a fragment mass tolerance of 0.01 Da, and a precursor mass tolerance of 10 ppm were involved in the search parameters. In addition, peptides were screened for modifications such as carbamidomethyl cysteine, sulfenylated cysteine, oxidized methionine, as well as deamidated glutamine and asparagine. Mitochondrial annotations were matched with the MitoCarta3.0 data set (Rath et al., 2021), and TOM70 substrates were extracted and compared according to the previous publications (Morgenstern et al., 2021).

#### Lipidome Profiling by UHPLC−MS

For lipidomic analysis, wild-type and TMX2 KO cells were harvested and washed 3 times with pre-cooled PBS (1.25 × 10^7^ cells/sample). Next, the tubes with cell pellets were kept in liquid nitrogen for 2 min to stop the metabolism and stored in a −80 °C freezer (protection from light for further use). Lipids extraction was performed strictly following a modified Folch liquid-liquid extraction protocol (Zardini Buzatto et al., 2020). Briefly, samples were mixed with NovaMT LipidRep internal standard solution (a mixture of 15 deuterated lipid standards), dichloromethane and methanol. After homogenization and cleaning with water, samples were equilibrated for 10 min at R.T. and spun at 16,000 g for 10 min at 4°C. An organic layer was harvested and evaporated to dryness using a nitrogen blowdown evaporator. Dried samples were then resuspended in NovaMT MixB, vortexed for 1 min, and diluted with NovaMT MixA (Nova Medical Testing Inc., Edmonton, Canada). For quality control (QC), a pooled mixture randomly extracted from each sample was prepared.

LC-MS/MS was performed in positive and negative ionization to identify and quantify the extracted lipids. A Thermo Vanquish UHPLC linked to Bruker Impact II QTOF Mass Spectrometer was used, equipped with a Waters Acquity CSH C18 column, 1.7 µm (Thermo Fisher Scientific). MS/MS spectra were acquired for sample alignment, and identification was processed by NovaMT LipidScreener. MS/MS peaks were identified by a three-tier identification approach based on MS/MS spectral similarity, retention time, and accurate mass match (Tier 1: MS/MS match score ≥500, precursor m/z error ≤20.0 ppm, 5.0 mDa; Tier 2: MS/MS match score <500; precursor m/z error ≤20.0 ppm, 5.0 mDa; Tier 3: Mass match with m/z error ≤20.0 ppm and 5.0 mDa) (Easton et al., 2023). All compounds identified in these tiers were utilized for normalization and data analysis. The identified lipids were class-matched to the most similar internal standards (15 deuterated lipid standards) and median-normalized. Unidentified features were saved for further investigation but not employed for statistics. Statistical analysis was performed with NovaMT LipidScreener. Non-informative features were filtered out during data processing.

#### Drosophila mendelian ratios and lifespan

Progeny counts for control (*UAS-luciferase RNAi*) and experimental (*UAS-CG11007 RNAi*) flies and lifespan analysis was performed as previously described (Marcogliese et al., 2018). Briefly, for repo-Gal4 crosses, flies lacking the balancer: TM3, Sb1 had both the Gal4 and UAS-RNAi present making the expected genotype 50%. For at least three independent crosses the observed/expected number of flies was recorded. Because *nSyb-Gal4* is homozygous viable, 100% of F1 progeny are the appropriate genotype for *nSyb>luciferase RNAi* or *nSyb>CG11007 RNAi*. For lifespan, flies of the correct genotypes and separated by sex were aged at indicated temperatures. Flies were transferred to new vials every 3 days. Flies were assessed for lethality every day.

#### Drosophila behavior

Both climbing (negative geotaxis) and mechanical seizure induction (bang-sensitivity) were performed as previously described (Marcogliese et al., 2022). Briefly, 24 h before testing, flies are individually housed in vials with 1 mL of food. On the day of testing, flies are transferred to a clean empty vial and allowed to habituate for 1 min. After habituation, the vial is tapped on the bench to induce negative geotaxis behavior and the fly the time to climb 7 cm is assessed. Flies that failed to cross the 7 cm line were given a score of 30. Following negative geotaxis, the bang-sensitivity assay was performed by vortexing flies for 15 s at maximum speed to determine if there are any paralysis or spastic behavior after vortexing. The researcher is blinded to genotype and drug. Total n derives from three independent crosses.

#### Data analysis

Statistical analyses were performed with GraphPad Prism^®^ 9.0 software (US). Values are expressed as mean ± SD. The Shapiro-Wilk test and QQ plots were performed to check data normality. For normally distributed data, unpaired t-test, one-way, and two-way analysis of variance (ANOVA) were performed. For non-normally distributed data, the Mann-Whitney U test was used to compare differences between two groups, and the Kruskal-Wallis test was used for three or more independent groups. Statistical significance was determined as p < 0.05.

### KEY RESOURCES TABLE

See separate document.

---

**Figure S1.**
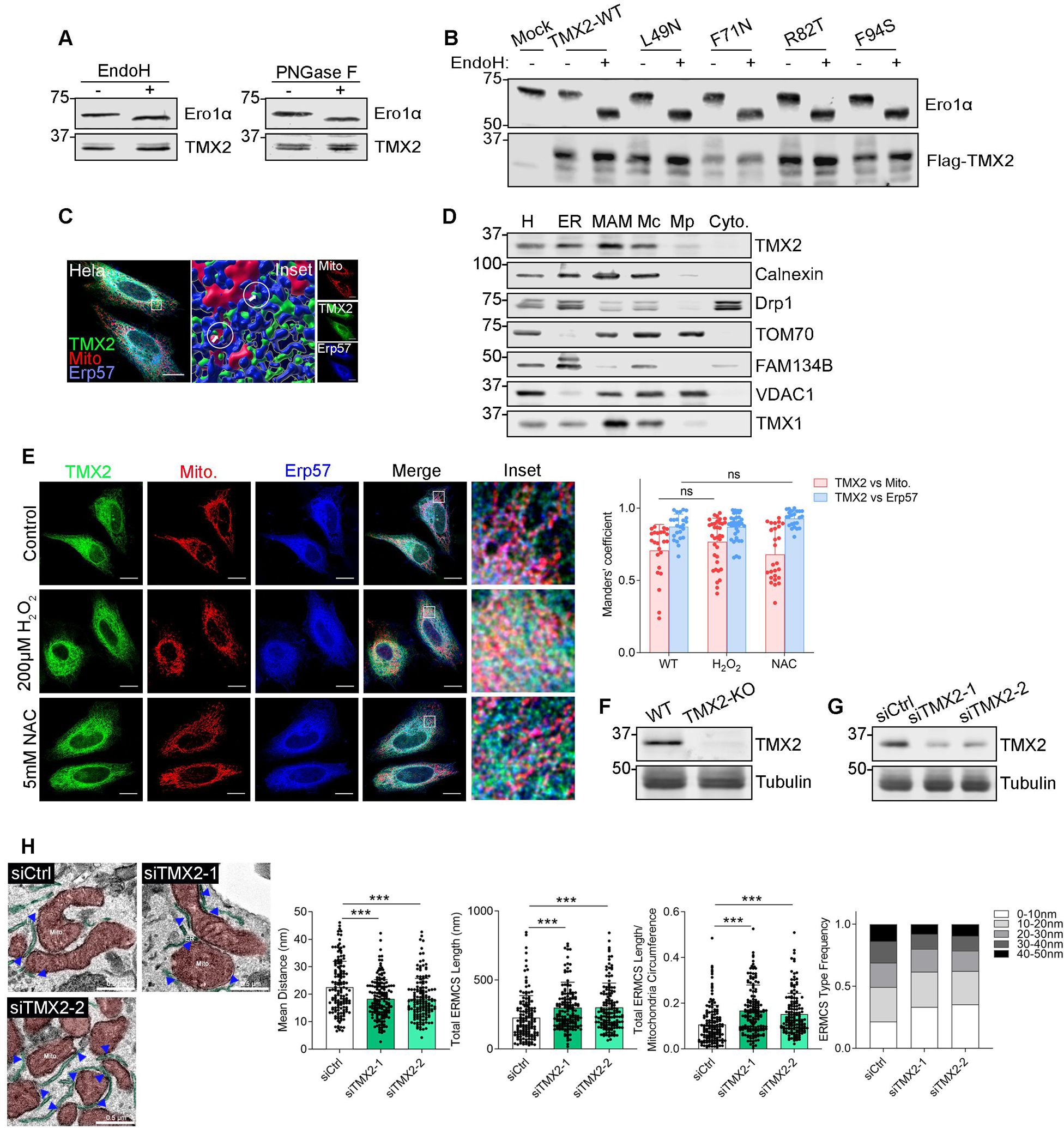
Supplementary information on basic TMX2 properties. (A) Glycosylation of endogenous TMX2. U251 MG cell lysates were treated with Endo H and PNGase F for 60 min at 37°C. Samples were then analyzed with Ero1ɑ and TMX2 antibodies. (B) Glycosylation scanning analysis. HeLa cells were transfected with the indicated artificial glycosylation mutants. Then, the glycosidase digestion assay was performed and the samples were analyzed with the Ero1ɑ and Flag antibodies. (C) 3D reconstruction of transfected Flag-tagged TMX2. HeLa cells were transfected with Flag-TMX2-wild-type for 24 h. Then immunofluorescence microscopy was performed, and TMX2 localization was observed after three-dimensional reconstruction (white arrow). Scale bar, 20 µm. (D) Percoll membrane fractionation of TMX2 in HeLa cells. Protein components of subcellular fractions prepared were analyzed by immunoblotting. Equal amounts were loaded and analyzed for the indicated ERMCS-regulatory proteins (H: homogenate; ER: endoplasmic reticulum; MAM: mitochondria-associated membrane; Mc: crude mitochondria; Mp: purified mitochondria; C: cytosol). (E) Redox dependence of TMX2 localization. After transfecting with Flag-TMX2-WT, HeLa cells were treated with 200 µM H_2_O_2_ for 20 min or 5 mM NAC for 1h. Mander’s coefficient analysis was utilized to measure the co-localization between TMX2 and MitoTracker^TM^ Red or Erp57 [n = 23-36 of 2 technical replicates for each group, statistical analysis was performed with two-way ANOVA (Sidak’s multiple comparisons test)]. Scale bar, 20 µm. (F, G) TMX2 expression levels after knockout and knock-down. TMX2-KO cells were generated with the CRISPR/Cas9 method using U251 MG cells. For TMX2 knock-down, U251 MG cells were transfected with RNAi (siTMX2) using Oligofectamine. Experiments were performed 48 h after transfection. The lysates were probed with a TMX2 antibody, and γ-tubulin served as a loading control. (H) ERMCS analysis in U251 MG TMX2 control and knock-down cells. Transmission electronic microscopy was performed with TMX2-KD cells. Mitochondria were labeled in red, and ER was labeled in green. Mitochondria circumference, average ER-mitochondria distance, and total ERMCS length were measured and analyzed [n = 69-166 of 3 technical replicates for each group, *p < 0.05, **p < 0.01, and ***p < 0.001 by Kruskal-Wallis test (Dunn’s multiple comparisons test)]. The ERMCS were classified and quantified by the mean distance. Scale bar, 0.5 µm.

**Figure S2.**
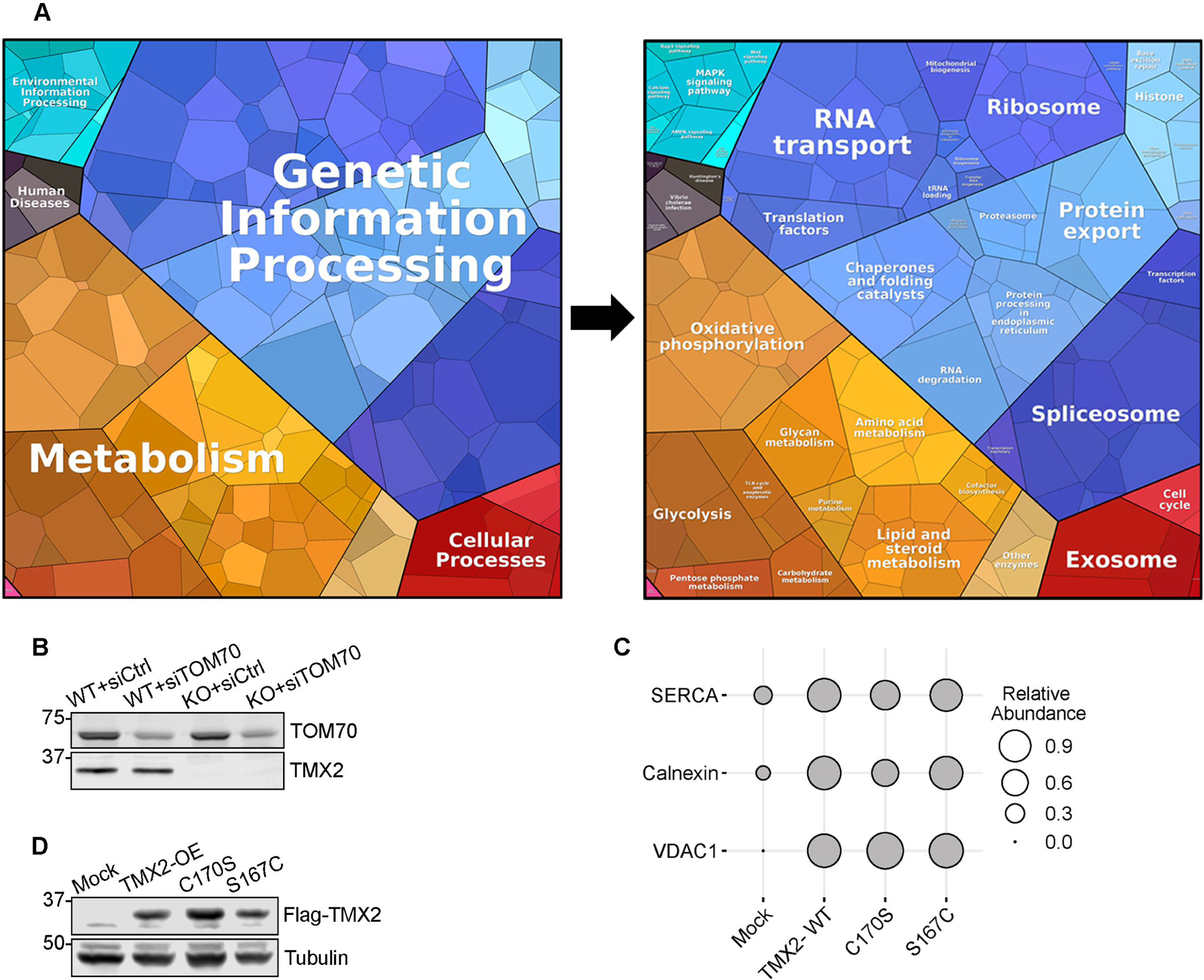
Supplementary information on TMX2 interactome. (A) Pathway analysis of the TMX2 interactome. Interacting proteins were grouped using the application Proteomap (http://bionic-vis.biologie.uni-greifswald.de/). Functionally related proteins are grouped in similarly colored polygons, and the area size reflects protein abundance. (B) TOM70 expression levels upon knock-down. U251 MG wild-type and TMX2-KO cells were transfected with TOM70 RNAi for 48 h, and then the cell lysates were subjected to the Western blot analysis using the indicated antibodies. (C) Analysis of additional ER and mitochondrial interactors of wild-type and mutant TMX2. Indicated TMX2 constructs (WT, C170S, and S167S mutants) were transfected and immunoprecipitated. The relative abundance of interacting partners with TMX2-wild-type and its mutants from the MS analysis was plotted as a bubble plot. Protein abundance was quantified with peak areas, which were normalized to TMX2-WT. (D) Expression levels of transfected wild-type and mutant TMX2 in HEK293 cells. The cell lysates extracted from the stable cell lines were subjected to the Western blot analysis using the indicated antibodies.

**Figure S3.**
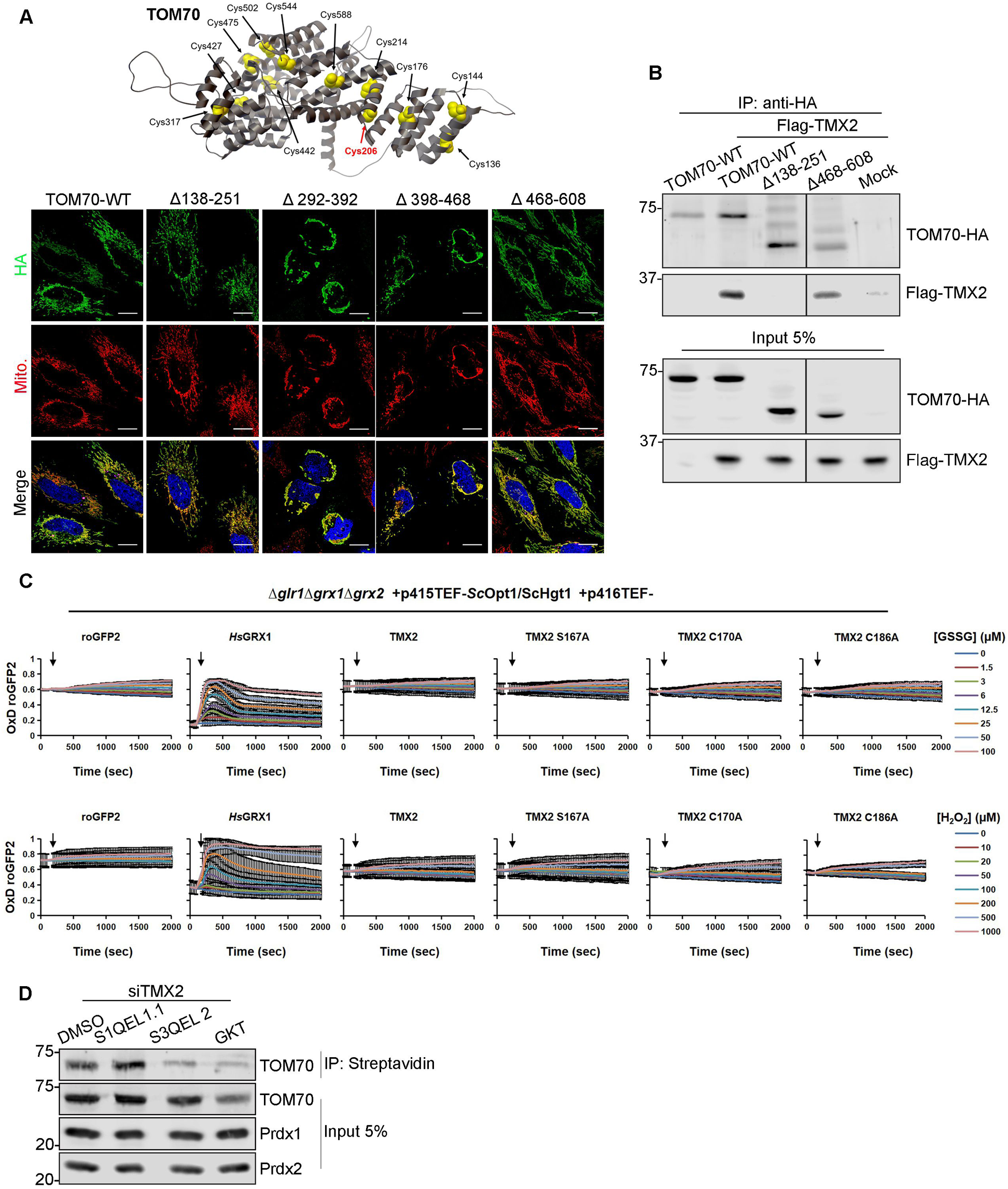
Supplementary data on TMX2-TOM70 interaction and effects on redox regulation. (A) Expression of TOM70 deletion mutants. HeLa cells were transfected with HA-tagged TOM70 truncations within the cytosolic domain (top), and the immunofluorescence microscopy was conducted 24 h after transfection. Scale bar, 20 µm. (B) Interaction of TMX2 with TOM70 deletion mutants. The co-immunoprecipitation assay was performed using HEK293 cells co-transfected with Flag-TMX2-wild-type and HA-tag TOM70 truncations for 24 h. Proteins associated with Flag-TMX2 were probed with a HA antibody. (C) Assay to measure TMX2 oxidative activity. Cytosolic roGFP2-ScGrx1 oxidation was measured in YPH499 *Δglr1Δgrx1Δgrx2* cells (containing TMX2 plasmids) following the addition of exogenous GSSG or exogenous H_2_O_2_ at the indicated concentrations. (D) Origin of ROS mediating TOM70 sulfenylation in TMX2 knock-down cells. HEK293 cells were transfected with TMX2 RNAi for 48 h and treated with 15 µm S1QEL1.1 (Complex I inhibitor), 15 µm S3QEL 2 (Complex III inhibitor), and 5 µm GKT (NOX4 inhibitor) for 20 h. Then the DCP-Bio1 assay was performed as described.

**Figure S4.**
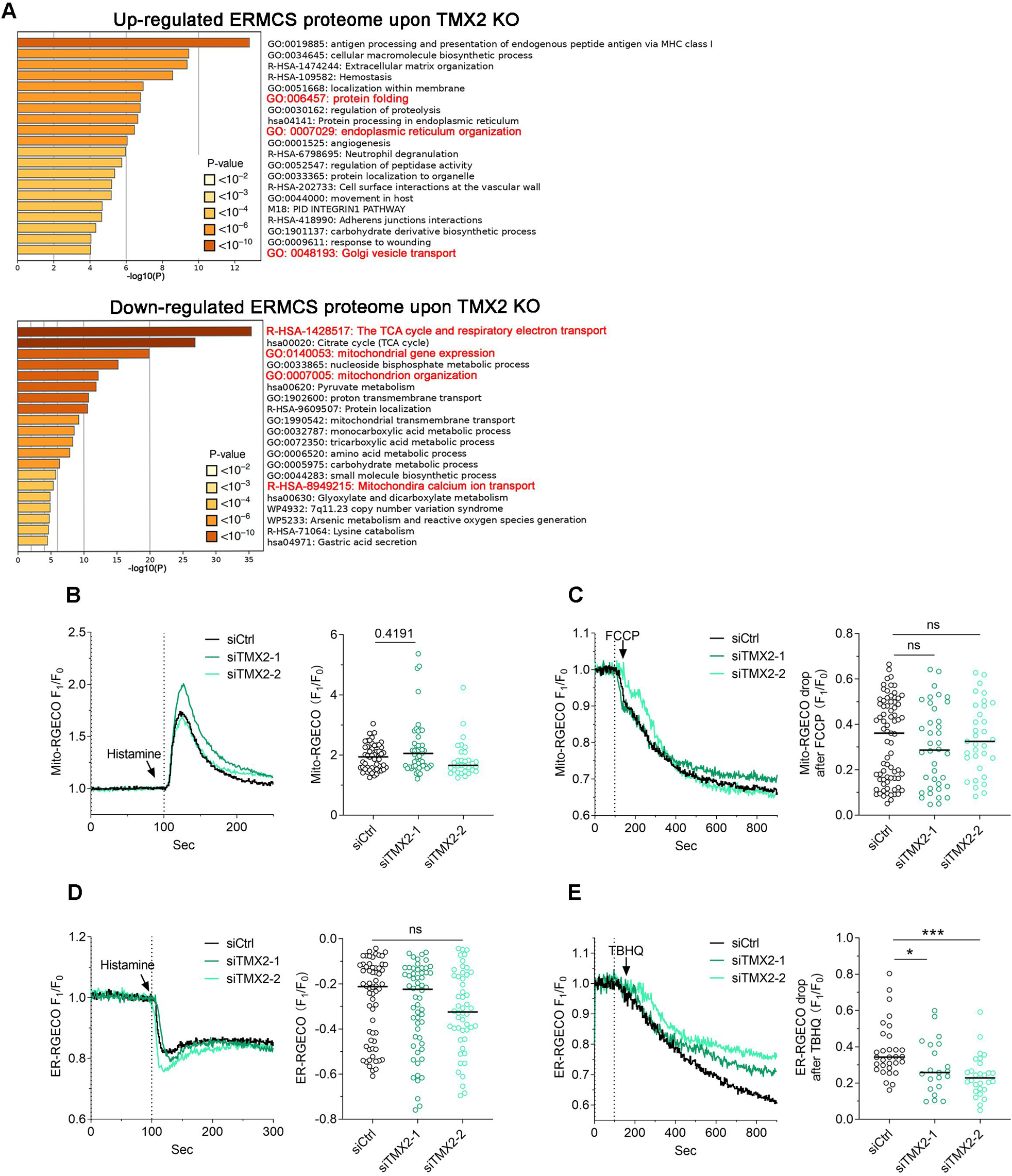
Supplementary data on ERMCS proteome and Ca^2+^ flux. (A) Pathway analysis of up- and down-regulated MAM proteome in U251 MG TMX2 knockout cells. Based on the MAM proteome data, pathway enrichment analysis was performed using the Metascape database. A bar chart showed the representative term clusters altered in the absence of TMX2, and the color represented the significance of the pathway enrichment. (B-E) Ca^2+^ responses from control and TMX2 knock-down cells as indicated. (B) U251 MG cells were transfected with indicated siRNA for 48 h and the plasmid encoding Mito-RGECO for 24 h. The Ca^2+^ signal was induced with 50 µm histamine. Relative fluorescence increases were recorded and quantified [n = 31-55 of 3 biological replicates for each group, statistics were performed via the Kruskal-Wallis test (Dunn’s multiple comparisons test)]. (C) U251 MG cells were transfected with indicated siRNA for 48 h and the plasmid encoding Mito-RGECO for 24 h. Then the cells were incubated with 10 µM FCCP to block mitochondrial Ca^2+^ import. Live-cell images were recorded and quantified [n = 33-75 of 2 biological replicates for each group, statistics were performed via the Kruskal-Wallis test (Dunn’s multiple comparisons test)]. (D) U251 MG cells were transfected with indicated siRNA for 48 h and the plasmid encoding ER-RGECO for 24 h. Then ER Ca^2+^ release under IP_3_Rs activation (50 µm histamine) was performed. The fluorescence drop was recorded and quantified [n = 50-62 of 2 biological replicates for each group, and statistical analysis was conducted by the Kruskal-Wallis test (Dunn’s multiple comparisons test)]. (E) U251 MG cells were transfected with indicated siRNA for 48 h and ER-RGECO for 24 h. Then cells were treated with 60 µM TBHQ, and the drop of the fluorescence signals was quantified (n = 21-31 of 2 biological replicates for each group, *p < 0.05 by Kruskal-Wallis test).

**Figure S5.**
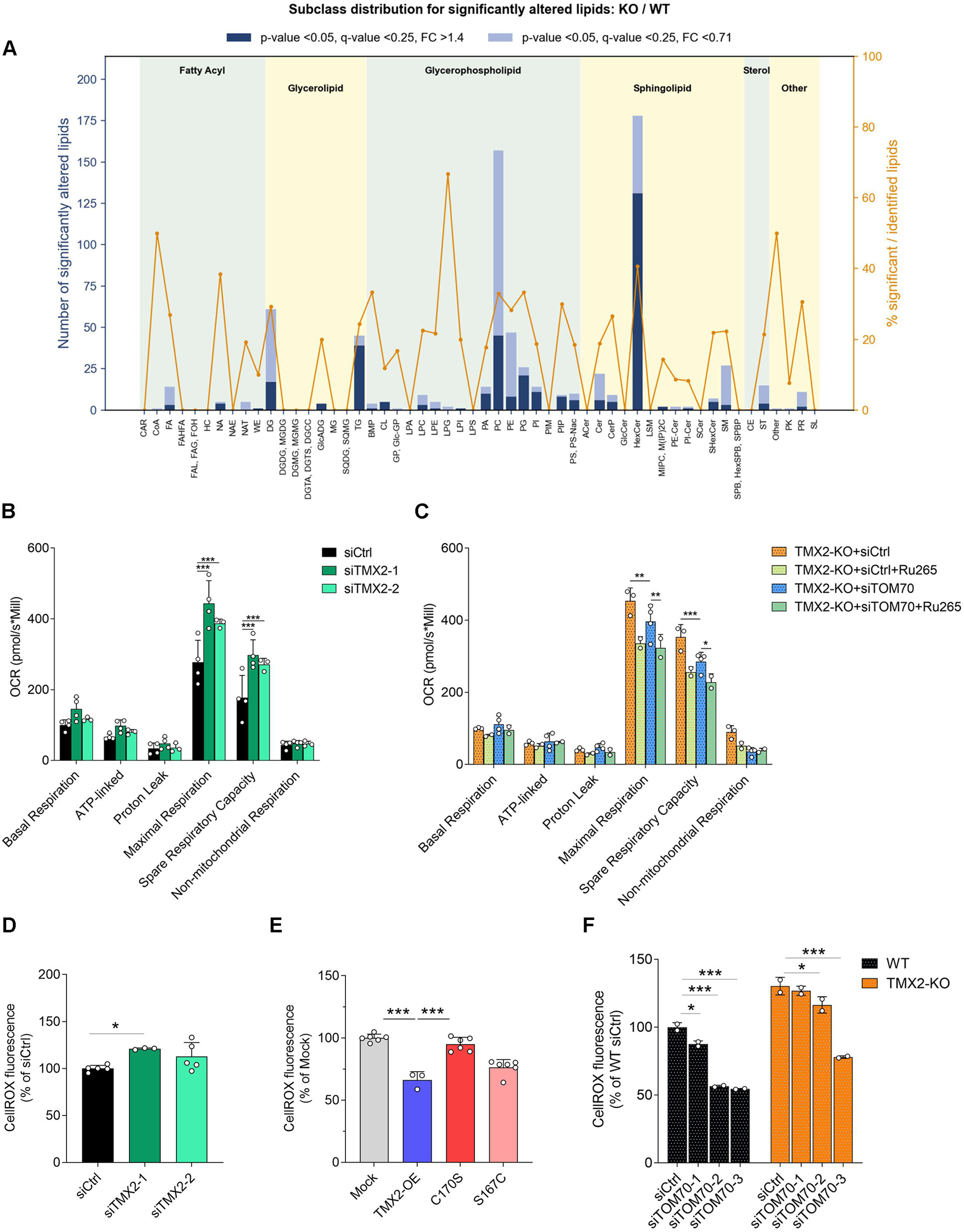
Supplementary data on cellular lipidomics and mitochondrial respiration. (A) Lipidome analysis from U251 MG wild-type and TMX2 knockout cells. The class distribution of the significantly altered lipid species (FC >1.40 or <0.71, p-value <0.05, and q-value <0.25) was shown by the bar graph. (B) Oxygen consumption rate in U251 MG wild-type and TMX2 knock-down cells. U251 MG cells were transfected with siRNA constructs for 48 h, and then mitochondria respiration measurements were performed [n = 3 biological replicates for each group, ***p < 0.001 by two-way ANOVA (Dunnett’s multiple comparisons test)]. (C) Ca^2+^ and TOM70 dependence of altered respiration in U-252 MG TMX2-KO cells. As indicated, cells were transfected with TOM70 RNAi for 48 h and treated with 25 µM of Ru265 (MCU inhibitor) overnight. Then, high-resolution respirometry was performed [n = 2-3 biological replicates for each group, *p < 0.05, **p < 0.01, and ***p < 0.001 by two-way ANOVA (Dunnett’s multiple comparisons test)]. (D-F) CellROX measurements as indicated. (D, E) U251 MG cells transfected with siRNA constructs for 48 h and U251 MG cells stably expressing TMX mutants were subjected to the measurements of cytosolic ROS production [n = 3 biological replicates for each group, *p < 0.05, and ***p < 0.001 by one-way ANOVA (Dunnett’s multiple comparisons test)]. (F) Wild-type and TMX2-KO cells were transfected with TOM70 RNAi for 48 h. Then, the cytosolic ROS production was monitored via flow cytometry (n = 2 biological replicates for each group, *p < 0.05, and ***p < 0.001 by two-way ANOVA (Dunnett’s multiple comparisons test)].

**Figure S6.**
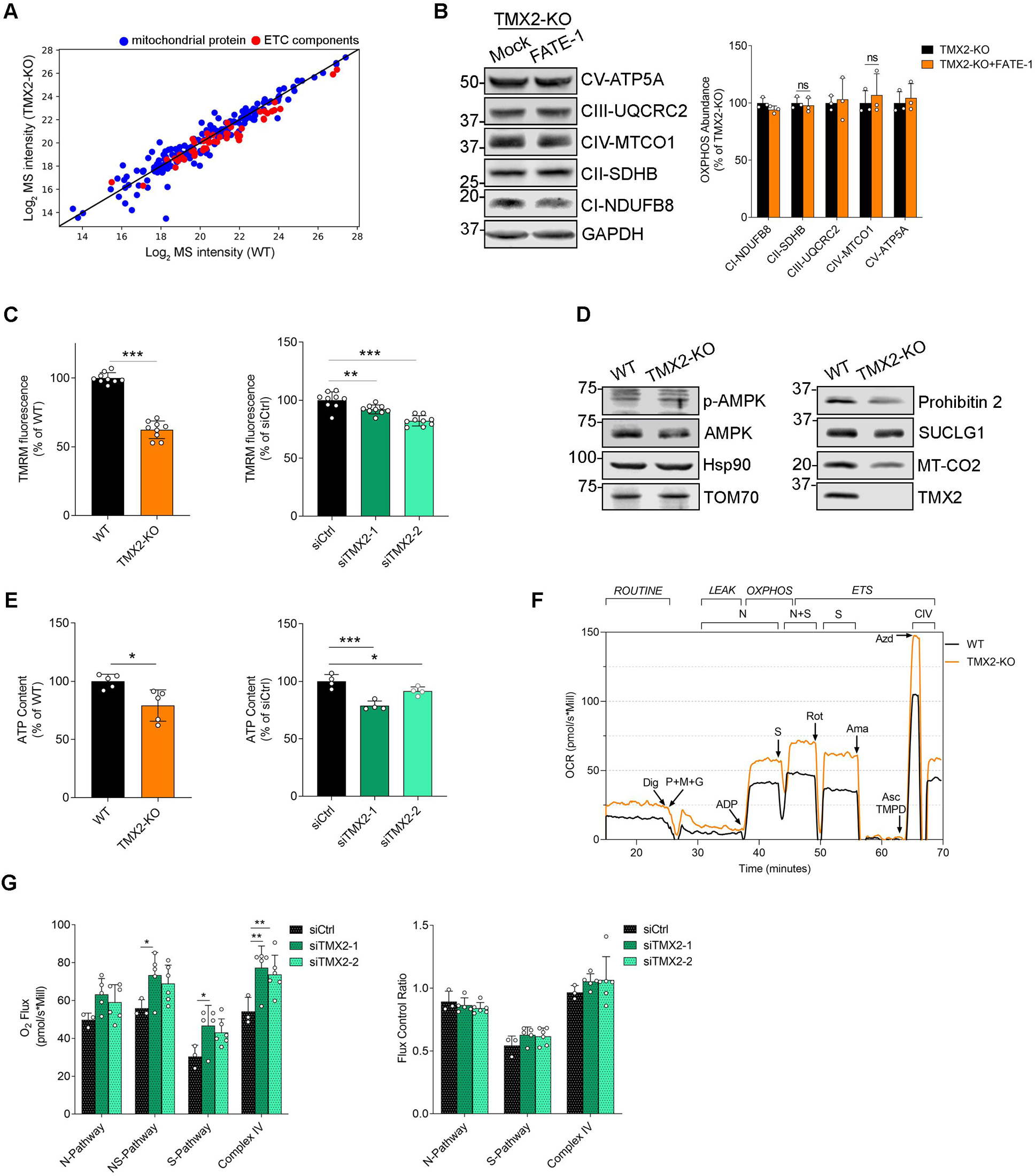
Supplementary information about mitochondrial proteome and function. (A) Correlation of protein abundance in the MS analysis of mitochondrial proteomes from wild-type and TMX2-KO cells. The ratio of mitochondrial protein abundance between wild-type and TMX2-KO cells was plotted in function of the protein abundance in wild-type cells. The ETC components were labeled in red dots. The lines indicate that the abundance of the protein between wild-type and TMX2-KO cells is the same. (B) Spacing reversal with FATE-1 does not revert U251 MG TMX2 knockcout proteome. U251 MG wild-type and TMX2-KO cells were transfected with FATE-1 for 48 h. Then, the cell lysates were collected and analyzed by Western blots with the indicated antibodies. (C) Mitochondrial membrane potential analysis. Wild-type and TMX2-KO cells, and U251 MG cells transfected with siRNA constructs for 48 h were used to measure mitochondrial membrane potential [n = 3-4 biological replicates for each group, **p < 0.01, and ***p < 0.001 by unpaired t-test or one-way ANOVA (Dunnett’s multiple comparisons test)]. (D) Additional lysate analysis from U251 MG wild-type and TMX2-KO cells. Cell lysates as indicated were collected and analyzed with Western blots with the indicated antibodies. (E) ATP content analysis. U251 MG wild-type and TMX2-KO cells, and U251 MG cells transfected with siRNA constructs for 48 h were used for the ATP measurements [n= 2-3 biological replicates for each group, *p < 0.05, and ***p < 0.001 by unpaired t-test or two-way ANOVA (Šídák’s multiple comparisons test)]. (F) Mitochondrial pathway analysis. Representative trace of the O_2_ flux in the digitonin-permeabilized wild-type and TMX2-KO cells. N-pathway (NADH-linked substrates entering the ETC via complex I) was monitored after adding pyruvate+malate+glutamase (P+M+G) and ADP. NS-pathway (electron flux convergence at Complexes I+II) was measured by the addition of succinate (S). S-pathway (succinate entering the RTC through Complex II) was tested after inhibiting Complex I with rotenone (Rot). Complex IV activity was measured in the presence of ascorbate (Asc) and tetramethylphenylenediamine (TMPD), transferring electrons to complex IV. The chemical background was subtracted after adding 100 mM sodium azide (Azd). Arrows indicate the time point of adding the substrates or inhibitors. (G) Quantification of mitochondrial pathway analysis. U251 MG cells were transfected with indicated siRNA for 48 h, and subjected to the mitochondrial respiration assay as described [n = 3-4 biological replicates for each group, *p < 0.05, and **p < 0.01 by two-way ANOVA (Dunnett’s multiple comparisons test)].

**Figure S7.**
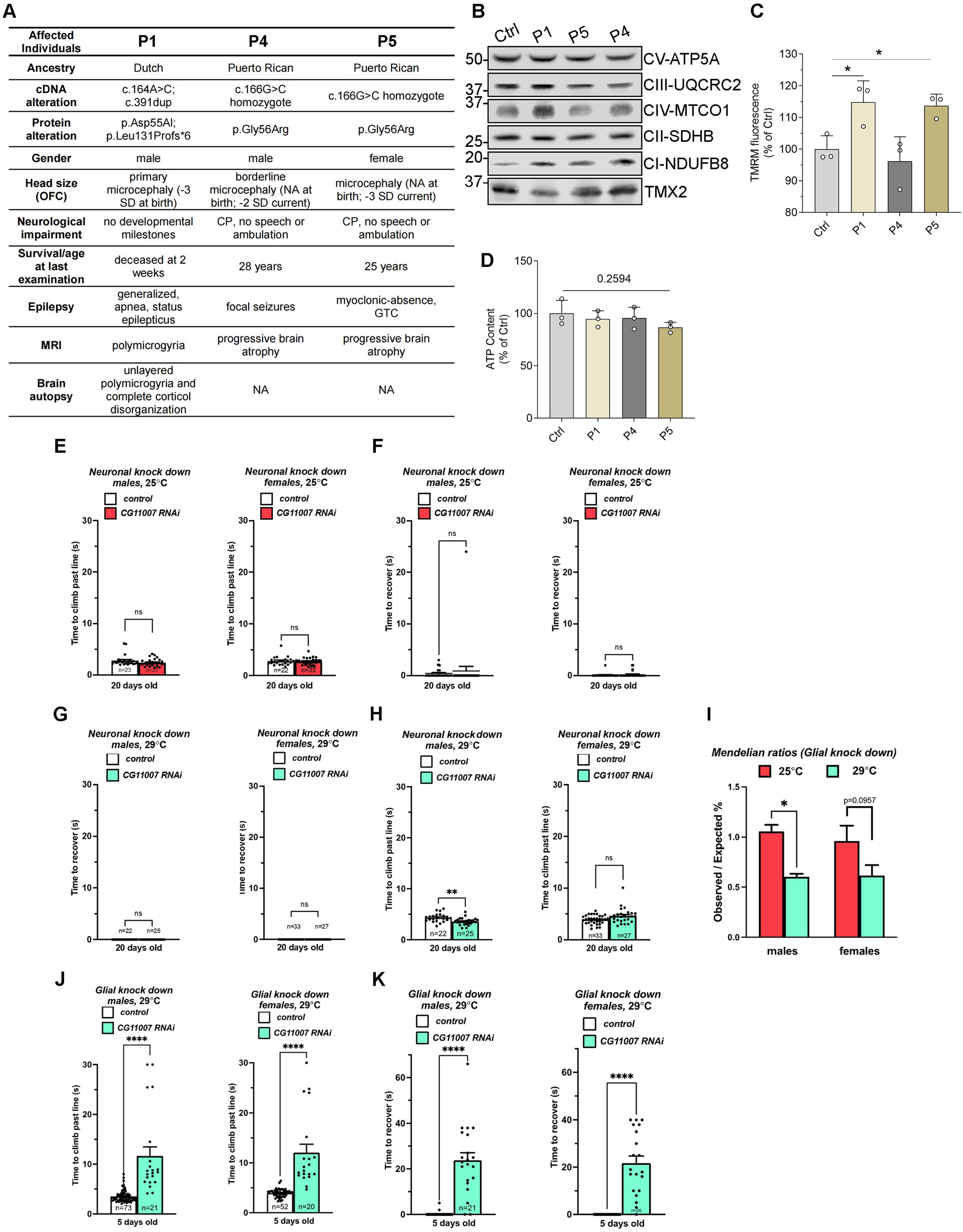
Supplementary data on *TMX2* mutant patient fibroblasts. (A) Summary of genetic variants and clinical characteristics in TMX2-linked patient fibroblasts. Adapted from (Vandervore et al., 2019). (B) Lysate analysis from patient cells. The cell lysates from patient fibroblasts were analyzed and probed with the indicated antibodies. (C) Mitochondrial membrane potential analysis. TMRM signals of patient fibroblasts was measured via flow cytometry [n = 3 biological replicates for each group, *p < 0.05 by one-way ANOVA (Dunnett’s multiple comparisons test)]. (D) ATP content analysis. The ATP measurements were performed with patient fibroblasts [n = 3 biological replicates for each group, statistical analyses were performed by one-way ANOVA (Dunnett’s multiple comparisons test)]. (E-H) Assays from neuronal *CG11007* knockdown. (E, F) Pan-neuronal knock-down of *CG11007* (nSyb-Gal4 > UAS-CG11007-RNAi) at 25°C does not cause any climbing deficits or mechanically induced seizures in male and female flies at 20 days post-eclosion (n = 3 biological replicates for each group, statistical analyses were performed via unpaired t-test). (G, H) Neuronal knock-down of CG11007 (nSyb-Gal4 > UAS-CG11007-RNAi) at 29°C leads to hyperkinetic climbing in males but fails to cause climbing deficits or mechanically-induced seizures in aged male and female flies at 20 days post-eclosion (n = 3 biological replicates for each group, **p < 0.01 by unpaired t-test). (I) Mendelian ratios of *CG11007* glial knockdowns. Increasing the incubation temperature to 29°C leads to significantly fewer repo-GAL4 > UAS-CG11007-RNAi that elcose from pupae than expected [n = 3 biological replicates for each group, *p < 0.05 by one-way ANOVA (Tukey post-hoc test)]. (J, K) High temperature analysis of glial *CG11007* knockdown. Pan-glial knock-down of *CG11007* (repo-GAL4 > UAS-CG11007-RNAi) at 29°C causes climbing deficits, mechanically induced seizures in young male and female flies at 5 days post-eclosion (n = 3 biological replicates for each group, ***p < 0.0001 by unpaired t-test).

## Notes

### Competing Interest Statement

The authors have declared no competing interest.

### Summary of Updates

The author name of Junsheng Chen was corrected.

